# LCK Regulates Homologous Recombination DNA Repair Identifying a New Target for Sensitizing PARP Inhibitors in HR Proficient Ovarian Cancer

**DOI:** 10.1101/2021.03.03.433791

**Authors:** Goutam Dey, Rashmi Bharti, Chad Braley, Ravi Alluri, Emily Esakov, Katie Crean-Tate, Keith McCrae, Amy Joehlin-Price, Peter G. Rose, Justin Lathia, Zihua Gong, Ofer Reizes

## Abstract

Poly-ADP Ribose Polymerase (PARP) targeted therapy is clinically approved for the treatment of homologous recombination (HR) repair deficient tumors. The remarkable success in treatment of HR repair deficient cancers has not translated to HR-proficient cancer. Our studies identify the mechanism of non-receptor lymphocyte-specific protein tyrosine kinase (LCK) in HR repair in endometrioid epithelial ovarian cancer (eEOC). LCK expression is induced and activation in the nucleus in response to DNA damage insult. LCK inhibition attenuates expression of RAD51, BRCA1, and BRCA2 proteins necessary for HR-mediated DNA repair, sufficient to suppress RAD51 foci formation, and augments γH2AX foci formation. Mechanistically, DNA damage leads to direct interaction of LCK with RAD51 and BRCA1 in a kinase dependent manner. Attenuation of LCK sensitized HR-proficient eEOC cells to PARP inhibitor in cell culture and pre-clinical mouse studies. These findings identify a new mechanism for expanding utility of PARP inhibitors in HR proficient ovarian cancer.

**Graphical Abstract:** 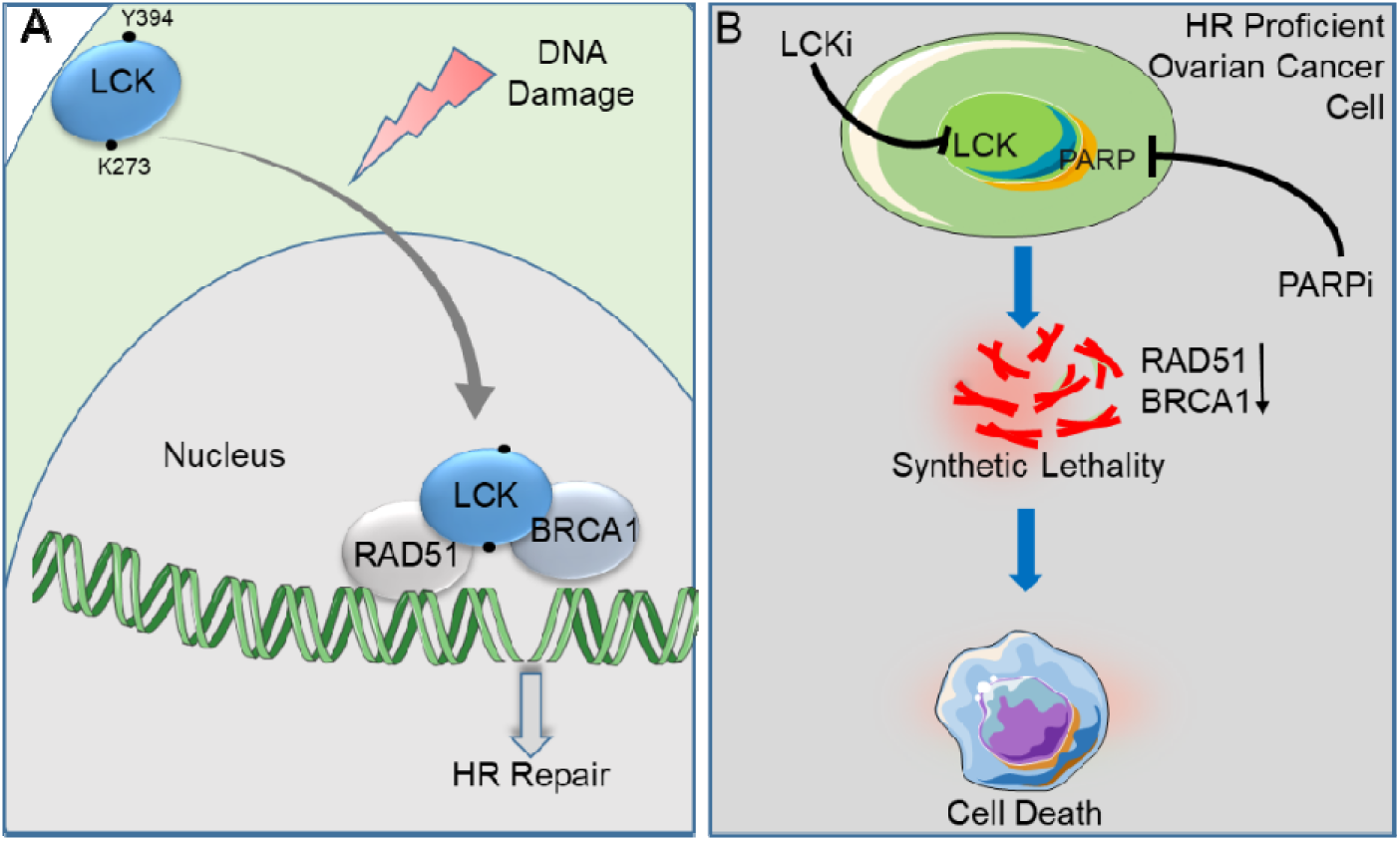

## Introduction

Epithelial ovarian cancer (EOC) is the second most common gynecologic malignancy in the United States, but the leading cause of gynecologic cancer death. It is estimated that in 2022, ∼19,880 women in the U.S. will be diagnosed with epithelial ovarian cancer (EOC) and 12,810 will die from their disease^1^. Poly-(ADP-ribose) polymerase inhibitors (PARPi) have emerged as new therapeutic options in the treatment of ovarian cancer^2–4^. Recent studies show that this treatment has prolonged median recurrence-free survival after primary therapy by more than 24 months^5^. While the benefit of PARPi is greatest in BRCA1/2-mutant or deficient tumors, those with HR deficiencies also experience a benefit from this therapy^5^. Conversely, PARPi and chemotherapy have so far shown limited efficacy in HR-proficient ovarian cancers. Further, platinum resistance is associated with HR proficiency in EOC^6, 7^. This limited efficacy of both platinum and PARPi therapy highlights an unmet clinical need in ovarian cancer patients.

Several strategies have been assessed to expand the utility of PARPi in HR-proficient cancers^2, 8–10^. RAD51, BRCA1, and BRCA2 are critical components of the HR repair complex. Studies have focused on disrupting this complex. Cyclin-dependent kinase (CDK) proteins demonstrably regulate the HR repair pathway in a lung cancer model^11^. Indeed, the CDK inhibitor dinaciclib is able to attenuate the expression of RAD51 and BRCA proteins resulting in the inhibition of HR repair capacity and potentiation of the pharmacological effect of PARPi^9^. However, there is no clinically approved drug for combination with PARPi for HR-proficient cancers.

Approximately 80% of endometrial cancers and 10% of ovarian cancers demonstrate endometrioid tumor histology (eEOC)^12^. A small but clinically significant proportion of eEOC display high-grade histology, advanced stage (FIGO stage III-IV), and a poor 5-year survival of 6-24%. These traits are similar to those of the more aggressive high-grade serous type of ovarian cancer ^13^. Moreover, somatic and/or germline mutations in HR genes occur in only a third of ovarian tumors, indicating the majority of eEOC are HR-proficient. Of note, eEOC show a considerably higher rate of resistance to platinum-based chemotherapy^14^ compared to serous carcinomas and do not commonly respond to targeted therapies such as PARP inhibitors.

We previously determined that intracellular, non-receptor tyrosine kinase, LCK regulates genes implicated in DNA repair machinery in eEOC^15^. We also demonstrated the pharmacologic inhibition of LCK attenuated expression of homologous recombination DNA damage repair genes leading to sensitization of eEOC cells to cisplatin^16^. In contrast, increased expression of LCK led to upregulation of DNA damage-repair genes and increased resistance to cisplatin. As LCK modulates RAD51, BRCA1, and BRCA2 expression, we hypothesized that blocking LCK expression or inhibiting kinase activity would sensitizes eEOC to PARPi. Here, we elucidate the mechanism of LCK in regulating HR DNA damage repair and a therapeutic approach to sensitize HR-proficient eEOC to PARP inhibitors.

## Results

### Regulation of Homologous Recombination (HR) DNA repair protein expression by LCK in eEOC

We tested whether LCK inhibition is sufficient to inhibit HR DNA repair genes RAD51, BRCA1, and BRCA2 at the protein level. We inhibited LCK expression using shRNA and CRISPR in the CP70 and SKOV3 cell lines, both of which are cisplatin resistant and known HR-proficient eEOC cells. Cells were transduced with lentivirus containing shRNA control (shCon) or LCK-targeted shRNA (KD1, KD2). Additionally, we generated LCK knock-out (KO) CP70 cells via CRISPR/Cas9. LCK inhibition was confirmed by immunoblotting followed by analysis of expression of BRCA1, BRCA2, and RAD51 via western blot analysis. In CP70 cells, we observed that KD1, KD2, and KO displayed attenuated protein expression of BRCA1, BRCA2 and RAD51 when compared to shCon (Fig. 1A, and Supplementary Fig. S1A and C). The protein expression levels of BRCA1, BRCA2, and RAD51 were similarly attenuated in LCK knock-down SKOV3 cells (Fig. 1A, and Supplementary Fig. S1B).

**Fig 1:**
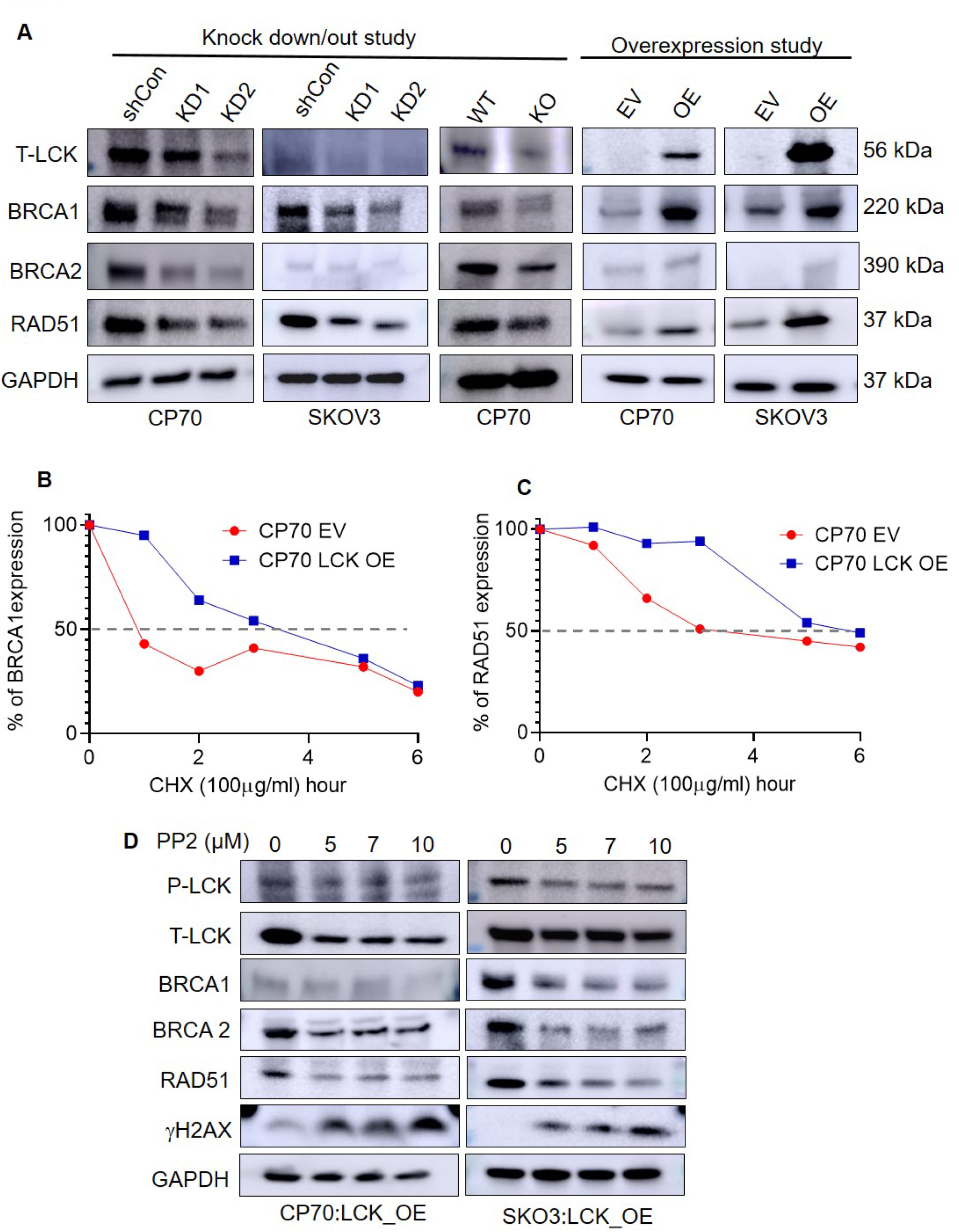
LCK modulates expression of HR repair proteins. **(A)** Western blot of CP70 and SKOV3 cells containing various lentiviral LCK KD, KO, and OE to determine effects on LCK, BRCA1, BRCA2 and RAD51 expression. **(B and C)** CP70 EV and CP70 LCK OE cells were treated with cycloheximide in a time dependent manner. Then, immunoblot analysis was performed to evaluate the expression of RAD51 and BRCA1 proteins. Half-lives were determined from digitized images. **(D)** Western blot analysis of CP70 and SKOV3 LCK OE cells treated with PP2 in a dose dependent manner for 48hrs, demonstrating the effects of a pharmacological inhibitor of LCK on P-LCK, T-LCK, BRCA1, BRCA2, RAD51 and γH2AX protein expression.

In complementary studies, we tested whether LCK overexpression would increase RAD51, BRCA1, and BRCA2 protein expression in eEOC. LCK overexpression led to induction of RAD51, BRCA1, and BRCA2 protein expression in CP70 and SKOV3 cells (Fig. 1A, and Supplementary Fig. S1D and E). To test the hypothesis that LCK increases the stability of BRCA1 and RAD51 proteins, empty vector (EV) and OE cells were treated with cycloheximide and harvested at various time-points to assess protein expression levels (Fig.1B, C, Supplementary Fig. S1F). The half-lives of BRCA1 and RAD51 were significantly increased in OE when compared to control. Half-lives of RAD51 in CP70 EV and OE cells treated with cycloheximide were 3.2 and 5.6 h, respectively. Half-lives of BRCA1 in CP70 EV and OE cells treated with cycloheximide were 54 min and 3.4 h, respectively. These studies indicate that LCK is sufficient to regulate BRCA1 and RAD51 protein expression via protein stabilization.

To test whether pharmacologic inhibition of LCK can attenuate expression of DNA damage repair proteins, we used PP2, a cell-permeable, small-molecule inhibitor of LCK kinase^17, 18^. We tested the efficacy of PP2 in CP70 and SKOV3 OE cells. PP2 attenuated pLCK at Y394, the autophosphorylation site of LCK in these cells (Fig. 1D, and Supplementary Fig. S1G and H). γH2AX, a marker of DNA damage and replication stress, was elevated by PP2 treatment. This is indicative of either increased damage or reduced repair of DNA damage due to attenuation of BRCA1 and RAD51 expression. LCK inhibition also attenuates expression of RAD51, BRCA1, and BRCA2 in parental CP70 and SKOV3 as well as in the CRL1978 clear-cell EOC cell line (Supplementary Fig. S2A).

### LCK inhibition attenuates HR DNA damage repair in eEOC

The inhibition of DNA damage repair genes led us to test whether LCK inhibition impairs HR-dependent DNA repair. We utilized the DR-GFP reporter assay established in U2OS cells to measure repair efficiency^19^ (Fig. 2A). U2OS cells with/without DR-GFP reporter system express endogenous LCK protein expression as shown by western blot analysis (Supplementary Fig. S2B). U2OS cells treated with PP2 leads to a dose-dependent reduction of the GFP-positive cell population when compared to DMSO-treated cell population, indicating reduced DNA repair as a consequence of LCK inhibition (Fig. 2B). Likewise, shRNA silencing of LCK led to a reduction in the GFP-positive population compared to shCon transduced cells (Fig. 2B). This indicates that LCK inhibition attenuates HR repair efficiency in cancer cells.

**Fig 2:**
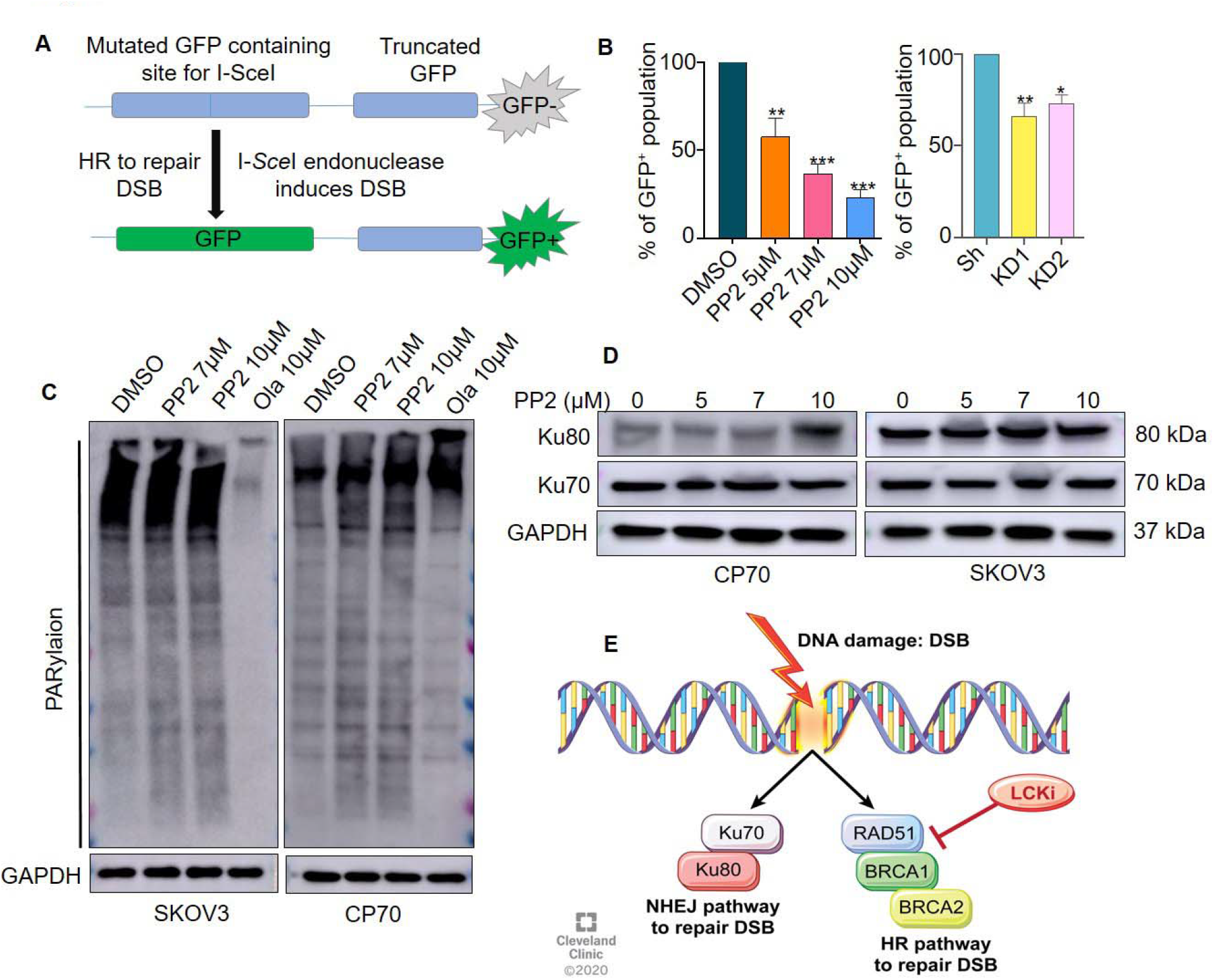
LCK inhibition attenuates the HR repair pathway in eEOC cells. **(A)** Schematic of DNA repair assay in U2OS osteosarcoma cells stably transduced with the DR-GFP reporter system. This reporter system contains upstream gene-encoding mutated GFP, and downstream truncated GFP. Transfection of I-SceI endonuclease induces double strand breaks in the upstream gene. Following efficient HR repair, GFP expression is restored and can be utilized to indicate HR-efficient cells for quantification. **(B)** U2OS cells were treated with varying concentrations of the LCKi PP2 for 48h or transfected with Sh Con, LCK KD1 or KD2 for 24h and incubated in serum enriched medium for another 24h. Cells were then subjected to DR-GFP assay. **(C)** CP70 and SKOV3 cells were treated with PP2 or Olaparib for 48h and immunoblot experiment was performed to check PARylation. (**D**) CP70, and SKOV3 cells were treated with increasing concentrations of PP2 for 48h and cells were harvested, lysed, and immunoblotted for Ku70 and Ku80 protein expression. GAPDH was used as loading control. **(E)** Schematic model summarizing LCK inhibition specificity for HR DNA repair. (Unpaired t test, p*<0.05, p**<0.01, p***<0.001).

DNA damage leads to activation of several repair pathways including PARP, HR, and NHEJ ^20^. As our studies indicated LCK inhibition attenuates HR repair proteins, we assessed LCK’s impact on the expression of alternative DNA repair pathways, including PARP and NHEJ, in CP70 and SKOV3 cells. The LCK inhibitor PP2 did not inhibit PARylation in CP70 and SKOV3 cells (Fig. 2C). The Ku80 and Ku70 proteins are a critical component of the NHEJ pathway^21^. After PP2 treatment, Ku80 protein expression was elevated in CP70, but not in SKOV3 (Fig. 2D). Furthermore, Ku70 protein expression was not changed in either CP70 or SKOV3 after PP2 exposure (Fig. 2D). In parallel, Ku70 and Ku80 expression levels in CRL1978 cells did not change following PP2 treatment (Supplementary Fig. S2C). These findings indicate that LCK inhibition targets HR repair proteins independent of induction of NHEJ repair mechanisms (Fig. 2E).

### DNA damage induces LCK dependent BRCA1 expression

We next assessed the effects of DNA damage on LCK expression and activation. DNA damage in ovarian cancer cells was induced using either etoposide, ultraviolet radiation, or methyl methanesulfonate (MMS). Dose-dependent treatment of CP70 cells with etoposide or MMS led to increased LCK protein expression (Fig. 3A and Supplementary Fig. S3A and B). Etoposide or MMS treatment in SKOV3 cells led to increased phosphorylation of LCK at pY394, while total levels of LCK protein remained unchanged (Supplementary Fig. S4A). Likewise, ultraviolet radiation of CP70 cells was sufficient to increase LCK phosphorylation (Supplementary Fig. S4B). BRCA1 and γH2AX expression was induced by etoposide or MMS, indicating increased DNA damage (Fig. 3A). Because DNA damage, particularly double strand breaks (DSB), and its repair machinery are concentrated in the nucleus^22^, we investigated the effects that DNA damage could induce on the accumulation of LCK in the nucleus. We found increased total LCK and pLCK in the nucleus of etoposide-treated cells (Fig. 3B and Supplementary Fig. S3C). Sub-cellular fractionation and immunofluorescence analysis both showed that pLCK was predominately localized to the nucleus of etoposide-treated cells (Fig. 3C). These findings were replicated in SKOV3 cells (Supplementary Fig. S4C). No pLCK was detectable in the nucleus of DMSO-treated cells (Fig. 3C). We next tested whether inhibition of LCK would be sufficient to block BRCA1 expression in etoposide-treated cells. Etoposide treatment in shCon cells showed increased BRCA1 protein expression, whereas this treatment attenuated BRCA1 expression in KD cells (Fig. 3D and Supplementary Fig. S3D). We repeated these studies in CRISPR/CAS9 KO cells and observed similar attenuation of BRCA1 in KO CP70 cells and no attenuation in parental cells (Fig. 3E and Supplementary Fig. S3E). These findings indicate that the induction of BRCA1 expression in response to DNA damage is disrupted by LCK inhibition.

**Fig 3:**
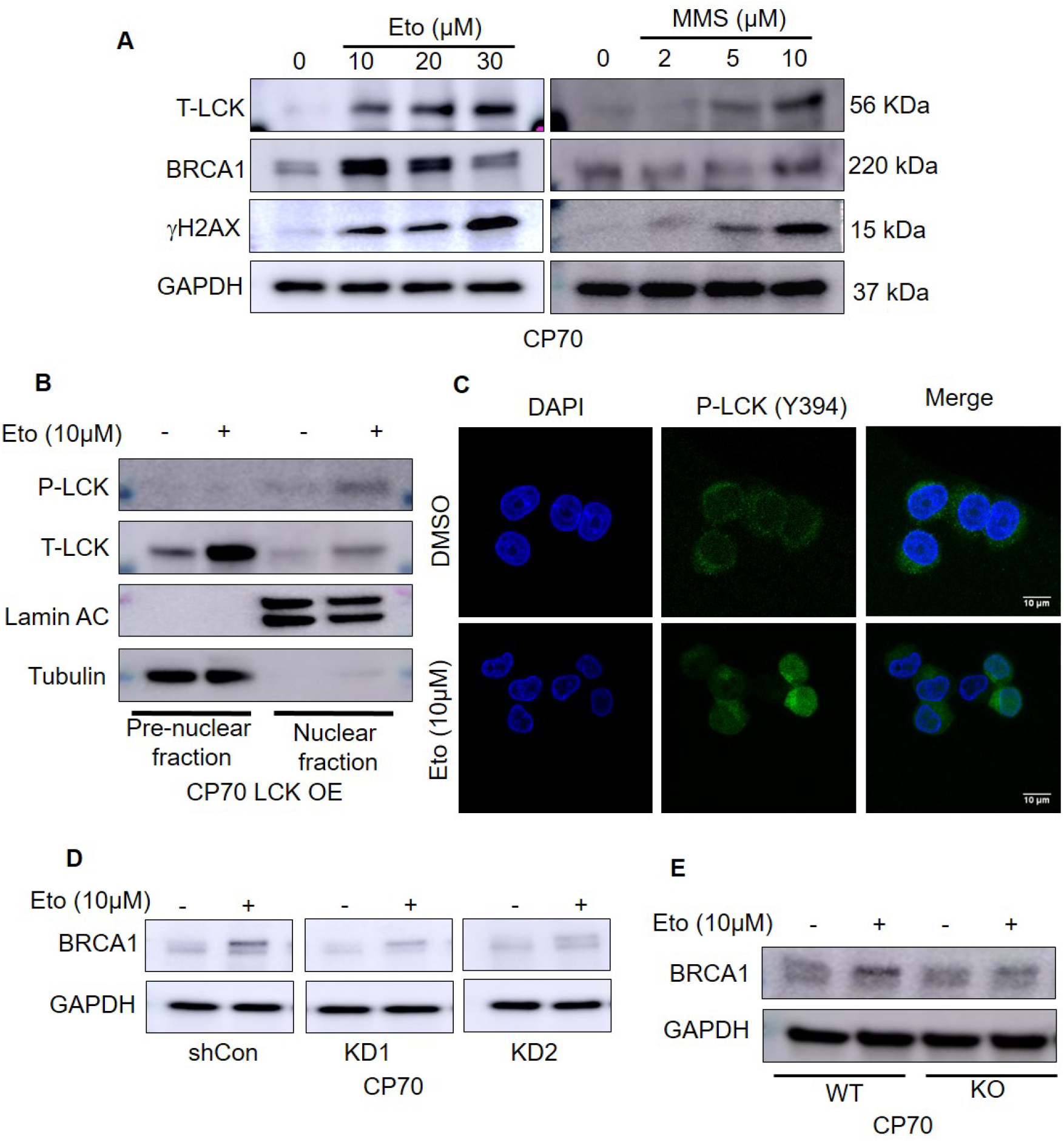
LCK is activated in the nucleus in response to DNA damage inhibition suppresses BRCA1 expression during DNA damage response. **(A)** Western blot analysis of T-LCK, BRCA1 and yH2AX expression following 24hrs etoposide or MMS treatment, followed by 24hrs recovery in CP70 cells. **(B)** Etoposide treated CP70 LCK OE cells were used for cytoplasmic and nuclear proteins extraction followed by western blot analysis. **(C)** Immunofluorescence study of etoposide treated CP70 cells. **(D)** Western blot analysis of etoposide/DMSO treated CP70 cells (Sh Con or LCK KD1 and KD2). **(E)** Western blot analysis of etoposide/DMSO treated CP70 cells (WT and LCK KO).

### LCK regulation of DNA double strand break repair

As γH2AX and RAD51 are markers of DNA damage and repair of DSB, we tested for foci formation in control and etoposide-treated cells. KO and OE CP70 (LCK overexpression in CRISPR/Cas9-background) cells were treated in presence or absence etoposide then subjected to immunofluorescence analysis to detect and quantify γH2AX foci (Fig. 4A, B and Supplementary Fig. S5A). In the absence of etoposide, no γH2AX foci formation was detected. Parental CP70 (WT) cells treated with etoposide led to increased γH2AX foci compared to DMSO treatment. KO cells treated with etoposide exhibited 4-5 fold increased foci formation compared to WT. Foci formation was nearly completely blocked in etoposide treated OE cells (Fig. 4A and B).

**Fig 4:**
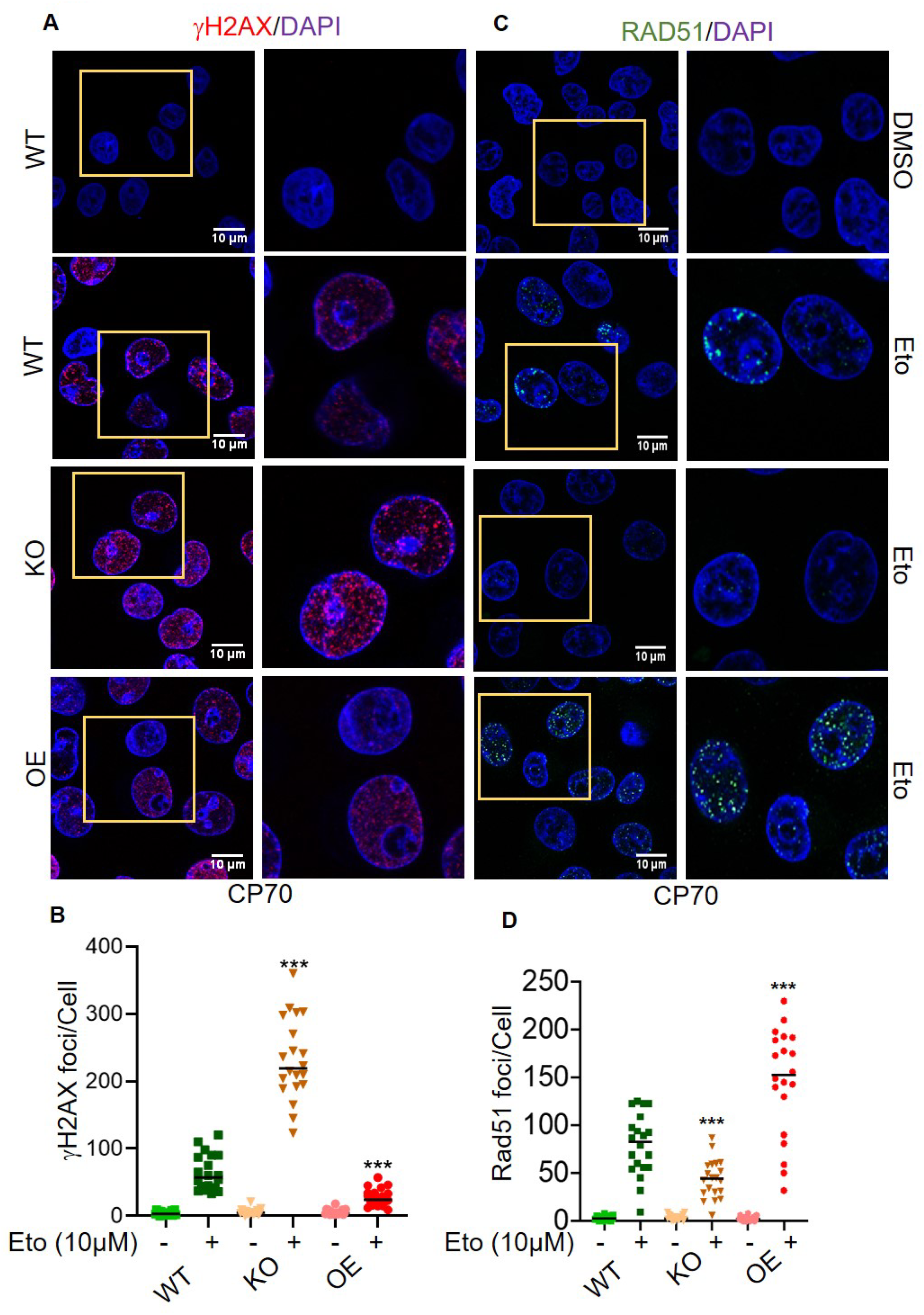
LCK regulates DNA damage and repair. **(A and B)** CP70 WT, LCK KO via CRISPR/Cas9 (KO), and LCK OE on CRISPR background (OE) cells were treated with DMSO/etoposide 10µM for 24h. Then cells were kept in drug-free media for another 24h. Immunofluorescence was performed to visualize γH2AX foci formation in different groups. Scale bar represents 10µm. γH2AX foci was counted by image J software and 20 cells were counted and plotted. **(C and D)** CP70 WT, LCK KO via CRISPR/Cas9 (KO), and LCK OE on CRISPR background (OE) cells were treated with DMSO/etoposide 10µM for 24h. Cells were incubated in drug free media for another 24h. Immunofluorescence was performed to visualize RAD51 foci formation in different groups. RAD51 foci in 20 cells were counted by image J software. Level of significance indicated on graph as determined by Graph pad Prism software (p*<0.05, p**<0.01, p***<0.001).

In parallel, we assessed RAD51 foci formation in KO and OE CP70 cells treated in the absence or presence of etoposide (Fig. 4C, D and Supplementary Fig. S5B). As with γH2AX, no RAD51 foci were observed in WT, KO, or OE cells treated with DMSO. In contrast, etoposide treatment led to a significant increase in RAD51 foci in WT CP70 cells that was significantly suppressed in KO cells (Fig. 4C, D, and Supplementary Fig. S5B). RAD51 foci formation was significantly increased in OE cells treated with etoposide. This data supports the conclusion that LCK can regulate HR repair during DNA damage response (DDR).

### LCK complexes with RAD51 and BRCA1 in response to DNA damage

As pLCK is localized in the nucleus in response to DNA damage and can induce BRCA1 and RAD51, we tested whether LCK directly interacts with RAD51 and BRCA1 in nuclear extracts. CP70 and SKOV3 cells were transduced with a myc-tagged LCK (Fig. 5A and Supplementary Fig. S6). We treated cells in the absence or presence of etoposide, isolated nuclei, and performed an IP (immunoprecipitation) assay with myc antibodies. In untreated cells, neither BRCA1 nor RAD51 co-precipitated with mycLCK. In contrast, etoposide treatment resulted in co-precipitation of RAD51 and BRCA1 with mycLCK (Fig. 5B and C). In parallel, RAD51 could co-immunoprecipitate with LCK from etoposide-treated OE SKOV3 cells. LCK and BRCA1 were detected in RAD51 immunoprecipitates (Fig. 5D). These findings indicate that in response to DNA damage, LCK interacts with a complex containing RAD51 and BRCA1 (Fig. 5E).

**Fig 5:**
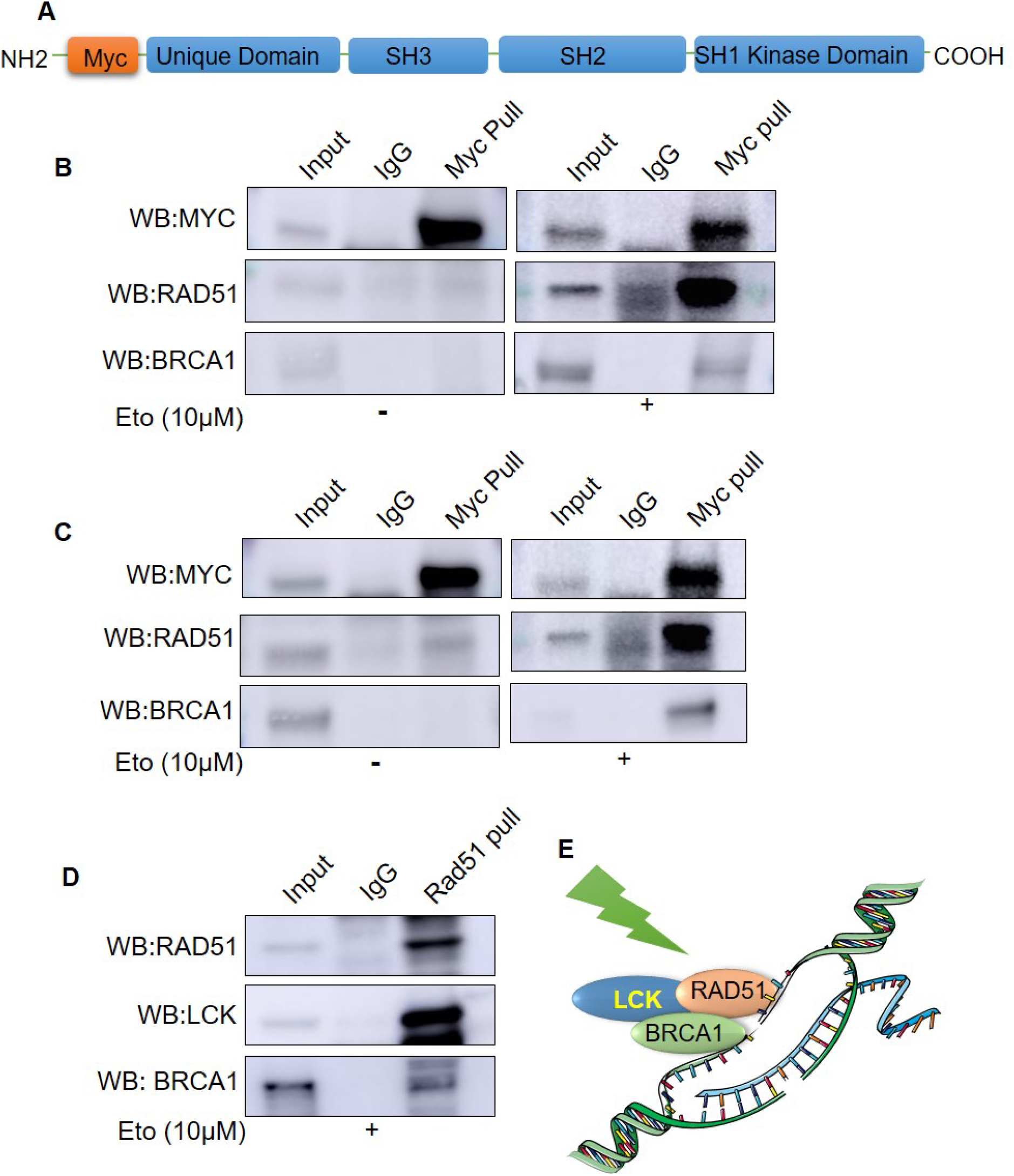
LCK interacts with RAD51 and BRCA1 during DNA damage response. **(A)** Schematic of Myc tagged LCK construct. **(B)** Myc tagged LCK expressing CP70 cells were treated with vehicle (DMSO) or etoposide. Cells were harvested, lysed, and nuclei were purified. Immunoprecipitation with Myc followed by immunoblotting for Myc, BRCA1, and RAD51. **(C)** Myc tagged LCK expressing SKOV3 cells were treated with vehicle or etoposide. Myc protein was pull down to determine the interaction of LCK with RAD51 and BRCA1 by co-immunoprecipitation study. **(D)** LCK overexpressing SKOV3 cells were treated with etoposide. RAD51 was pulled down from protein lysate. Then the expression of LCK and RAD51 was checked in pulled down protein sample by co-immunoprecipitation study. **(E)** Schematic of LCK binding partners during DNA damage response.

### LCK kinase activity is essential for HR repair

LCK interacts with BRCA1 and RAD51 in response to DNA damage, so we tested whether kinase activity and autophosphorylation of LCK is necessary for activity and DNA repair. We generated LCK mutants at lysine 273, which is necessary for kinase activity (K273R); tyrosine 394, an autophosphorylation and activation site (Y394F)^23^; and tyrosine 192, a SH2 adaptor protein binding site (Y192F)^24^ and transduced them into LCK KO CP70 cells (Fig. 6A). OE and Y192F mutants retained kinase activity, while K273R and Y394F mutants lacked kinase activity. We performed IP studies in etoposide-treated cells and determined that OE and Y192F cells were able to co-immunoprecipitate BRCA1 and RAD51, whereas K273R and Y394F failed to co-immunoprecipitated BRCA1 and RAD51 (Fig 6B). We next assessed γH2AX and RAD51 foci formation in the OE, K273R, Y192F, and Y394F transduced cells. OE and Y192F exhibited similar level of foci formation in response to etoposide (Fig. 6C, D, E, and F), whereas K273R and Y394F showed increased γH2AX foci and reduced RAD51 foci in etoposide treated cells (Fig. 6C, D, E, and F and Supplementary Fig. S7A, B). We further assayed the cells for etoposide sensitivity. We determined that naïve CP70 cells exhibited an IC50 of 5μM to etoposide. OE (IC50=10.03μM, 95% CI is 7.894 to 12.67) and Y192F (IC50=8.70μM, 95% CI is 7.044 to 10.72) exhibited increased resistance to etoposide, whereas K273R (IC50=2.28μM, 95% CI is 1.938 to 2.692) and Y394F (IC50=2.01μM, 95% CI is 1.655 to 2.455) showed increased sensitivity to etoposide (Fig. 6G). We performed a timed experiment with etoposide treated CP70 cells and determined that the extent of γH2AX foci was lower in OE and LCK Y192F mutant cells than in WT cells at 0 hours. Moreover, KO, Y394F, and K273R cells exhibited the highest number of γH2AX foci at all time points (Fig. 6H, and IF in Supplementary Fig. S8). These findings lead to the hypothesis that LCK kinase activity is essential for interaction with RAD51 and BRCA1 during DNA damage response and also facilitated HR repair.

**Fig 6:**
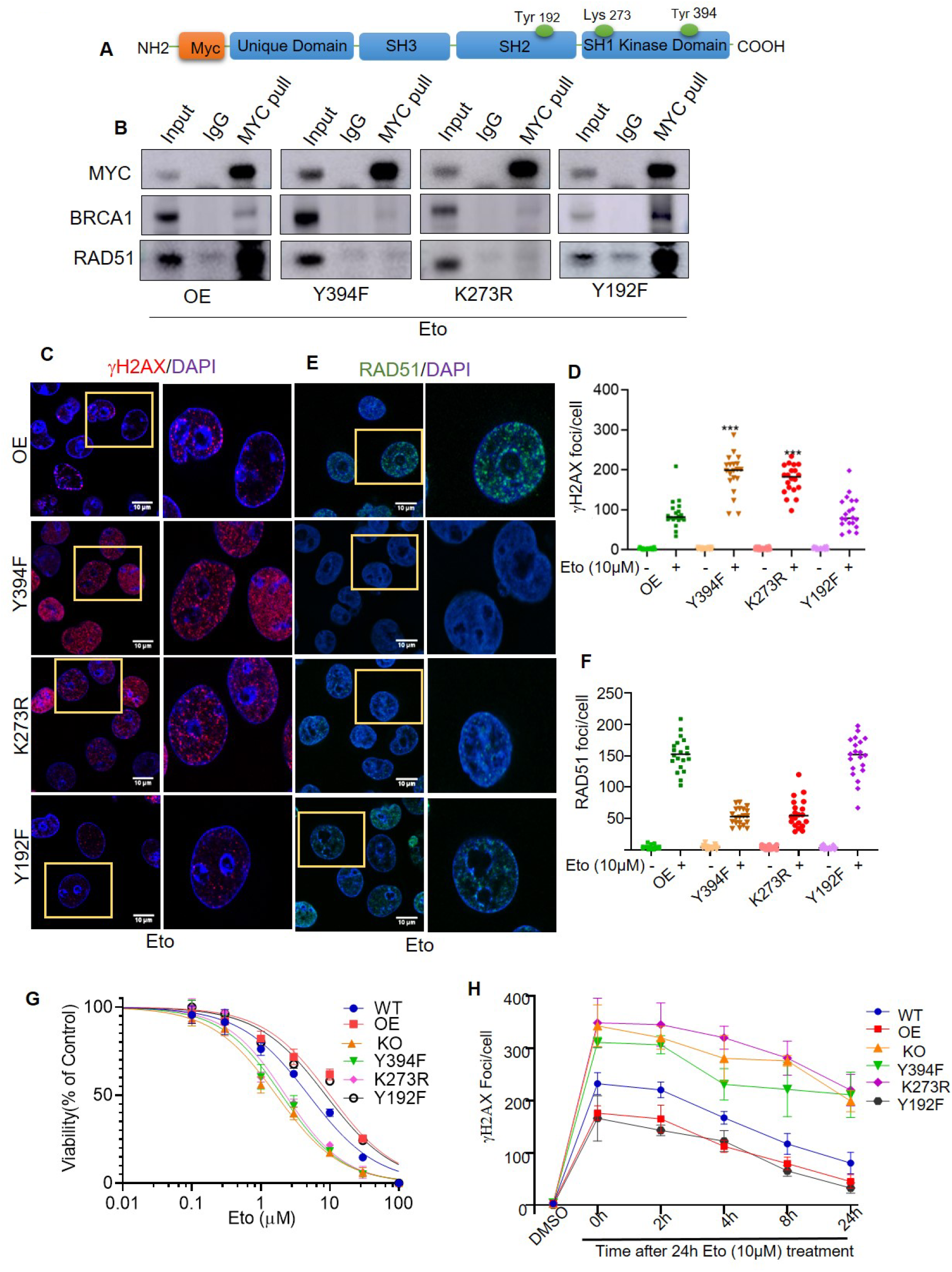
LCK kinase activity is essential for HR repair. **(A)** Structure of MYC labelled LCK construct. LCK Y394F, LCK K273R, LCK Y192F mutants were generated by site directed mutagenesis. These mutants were further introduced in CP70 cells having LCK KO background. **(B)** CP70 cells (MYC tagged LCK, LCK Y394F, LCK K273R, and LCK Y192F) were treated with etoposide/DMSO for 24h. Then cells were put in drug free media for 24h. Cells were collected and nuclear proteins were isolated. MYC antibodies were used for co-immunoprecipitation study to visualize the complex formation with RAD51 and BRCA1 proteins. **(C, E)** CP70 cells (LCK OE, LCK Y394F, LCK K273R, and LCK Y192F) were treated with etoposide for 24h. Cells were then kept in drug-free media for 24h. Immunofluorescence was performed to visualize γH2AX and RAD51 foci formation. **(D, F)** Quantification of γH2AX and RAD51 foci formation in CP70 cells by Image J software. **(G)** CP70 cells (WT, LCK OE, LCK KO, LCK Y394F, LCK K273R, and LCK Y192F) were treated with etoposide in a dose dependent manner for 48h. Then cell titer glow viability assay was performed to check cell viability. **(H)** CP70 cells (LCK, LCK Y394F, LCK K273R, and LCK Y192F in LCK knock out background) were grown on cover slips and treated with etoposide for 24h followed by incubation for 0, 2, 4, 8 and 24h. Cells were then subjected to immunofluorescence analysis to visualize γH2AX foci formation. Then γH2AX foci were counted and plotted to visualize the H2AX decay kinetics. p*<0.05, p**<0.01, p***<0.001 based on Graphpad Prism analysis.

### LCK inhibition augments PARPi induced DNA damage and genomic instability

We performed single cell gel electrophoresis (alkaline COMET) assay to quantify the extent of double and single strand DNA breaks by visualizing tail area^25^. CP70 and SKOV3 cells were incubated with either PP2, olaparib, or both. Treated cells were processed and stained with SYBR Gold to detect and measure the tail moment (Fig. 7A). PP2 and olaparib alone displayed a comparable increase in comet tails compared to DMSO (Fig. 7B). The combination of PP2 and olaparib induced a fourfold increase in comet tail area compared to monotherapy (Fig. 7B).

**Fig 7:**
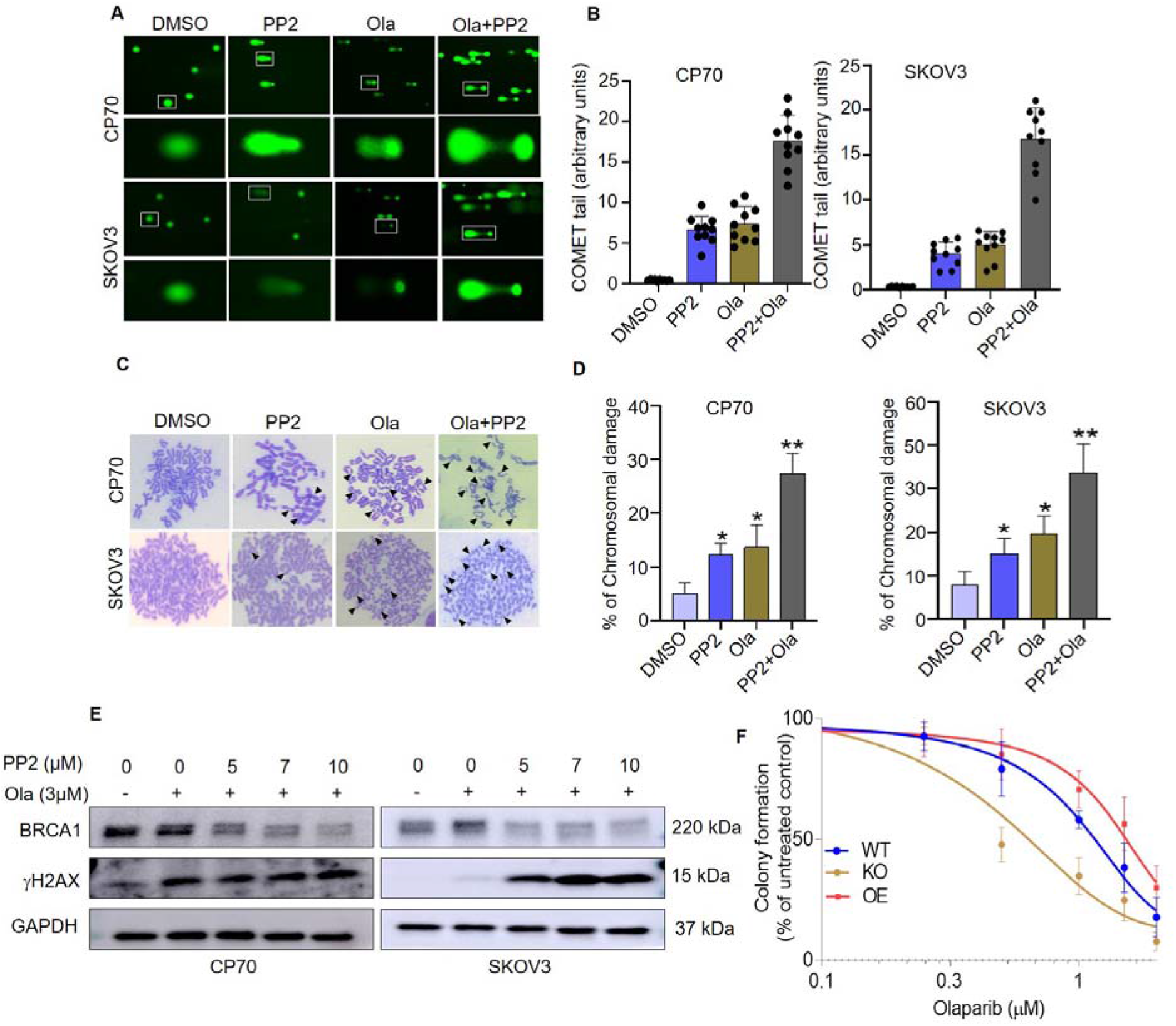
LCKi promotes genomic instability and augments PARPi induced genomic instability in ovarian cancer cells. **(A)** Single cell electrophoresis or COMET assay was used to validate and independently quantify the observed DNA damages. CP70 and SKOV3 cells were treated with PP2 5µM and/or Olaparib 3µM for 48h. Cells (1×10^5^ Cells/ml) were then collected and mixed with LMAgarose (1:10 V/V). Then, 50L of LMAgarose solution was put on COMET slide and subjected to single-cell gel electrophoresis. **(B)** Extent of DNA damage was estimated based on measurement of COMET tail area using Image J. **(C)** CP70 and SKOV3 cells treated with PP2 5µM and/or Olaparib 3µM for 48h then had the chromosomal aberration assay performed. The arrow indicates the presence of abnormalities in chromosomes including breaks, gaps, and radials. **(D)** Abnormities in chromosomes were quantified (Chromosomal break, gap, radial formation) by counting by visual observation. **(E)** CP70 and SKOV3 cells were treated with Olaparib and PP2 for 48h. Cells were harvested, lysed, and blotted for BRCA1 and γH2AX. **(F)** CP70 Parental (WT), KO, and OE (in KO background) cells were treated with Olaparib in dose dependent manner for 12 days. To identify colonies, plates were stained with crystal violet and images were captured. Colonies formed were counted and plotted as percentage of control formation in the gra. p*<0.05, p**<0.01, p***<0.001 based on Graph pad Prism analysis.

PARP inhibitors have been reported to induce genomic instability, leading to chromosomal aberration and DNA damage in cancer cells^26, 27^. Chromosomal damage can be detected by chromosomal breaks, gaps, and radial formations. We identified multiple breaks, gaps, and radial formation in PP2 and olaparib-treated cells (Fig. 7C). PP2 and olaparib displayed a comparable increase in chromosomal damage when compared to DMSO (Fig. 7D). The combination of PP2 and olaparib displayed increased chromosomal damage in both CP70 and SKOV3 cells (Fig. 7D).

Based on this analysis, we assessed whether the LCK inhibitor PP2 could synergize with a PARPi, olaparib, to augment the DNA damage response (Fig. 7E). Olaparib treatment led to an increase in BRCA1 expression and a detectable increase in γH2AX expression in SKOV3 cells (Fig. 7E). Co-treatment with PP2 was sufficient to suppress BRCA1 expression and significantly augment γH2AX expression in a dose-dependent manner (Fig. 7E). Our findings indicate LCK inhibition leads to HR deficiency. As proof of concept, we tested the impact of LCK silencing on the efficacy of olaparib in SKOV3 and CP70 cells via colony formation assay. Olaparib sensitivity was analyzed in parental (WT), KO, OE (Fig. 7F). We quantified colony formation and determined that sensitivity of CP70 to olaparib is greater in LCK KO than in parental CP70 cells (Fig. 7F, Supplementary Fig. S9). We determined that, in CP70 cells, olaparib resistance increased 3-fold in LCK OE compared to LCK KO. We replicated these findings by silencing with shRNA in CP70 and SKOV3 cells (Supplementary Fig. S10A and B). shCon, KD1, and KD2 cells were treated with various concentrations of olaparib and plated for colony formation. In CP70 cells, silencing LCK inhibited colony formation with greater efficiency in olaparib-treated cancer cells than in shCon treated cells (Supplementary Fig. S10A, B). Similarly, in SKOV3 cells, the number of colonies were significantly decreased after olaparib treatment in KD1 and KD2 cells as compared to shCon cells (Supplementary Fig. S10C, D). These findings support the hypothesis that olaparib has a higher efficacy in LCK-deficient cancer cells and indicate that LCK inhibition is sufficient to sensitize eEOC to PARPi.

### LCK inhibition potentiates therapeutic efficacy of PARPi in *in vivo*

To test whether LCK impacts olaparib efficacy in pre-clinical models of eEOC, we injected KO and OE CP70 cells into mice and once tumors were detected, we treated with 3 course of 5 day treatment of Olaparib (Fig. 8A). KO and OE CP70 exhibited nearly identical tumor growth in vehicle-treated mice (Fig. 8B and C). Olaparib treatment led to suppression of tumor growth in OE mice and to complete suppression of tumor growth in KO mice (Fig. 8B and C). These findings indicate that LCK inhibition potentiates olaparib synthetic lethality.

**Fig 8:**
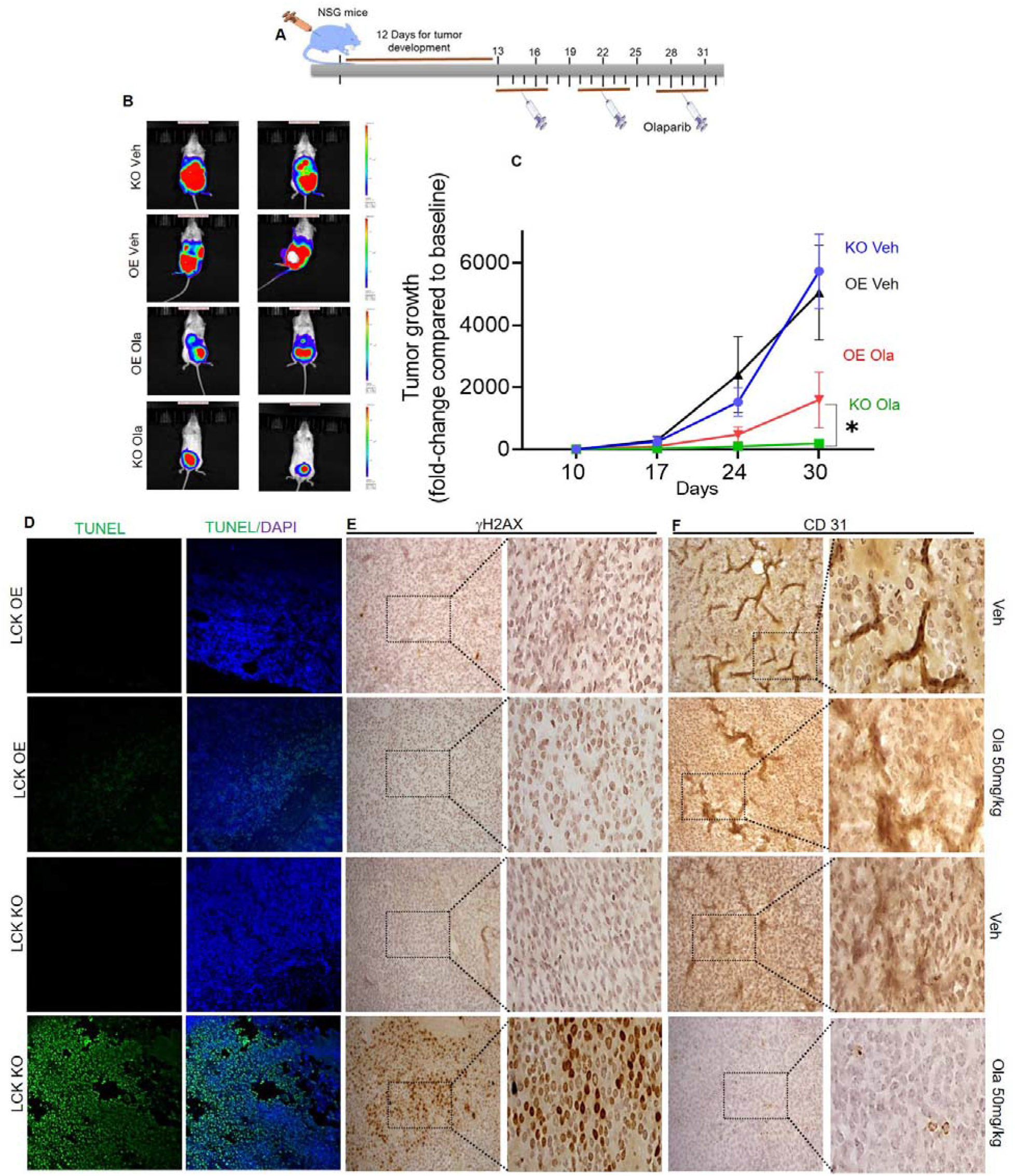
Disruption of LCK leads to inhibition ovarian tumor treated with Olaparib. **(A)** Schematic model of animal study. CP70 LCK KO and LCK OE cells were injected intraperitoneally in NSG mice. After 12 days, PARP inhibitor Olaparib i.p. (50mg/kg) was administered 5days/week. **(B)** After three weeks of Olaparib treatment, tumor volume was detected by IVIS imaging. **(C)** Tumor growth kinetics. **(D)** TUNEL assay to detect DNA fragmentation in tumor tissue sections. TUNEL positive cells were counted from five images and plotted in graph (Supplemental file S11 A). **(E)** IHC staining of γH2AX of tumor sections from different groups. γH2AX positive cells were counted from five images and plotted in graph (Supplemental file S11 B). **(F)** CD31 expression (Indicator of microvessel density and growth) of tumor sections from different group of mice. Microvessel density was counted from five images and plotted in graph (Supplemental file S11 C). Images are representative of two tumors from each cohort. We quantified the staining from 5 fields from each mouse. Images were captured at 20X magnification.

We performed a molecular analysis on tumor sections from OE and KO cells treated with and without olaparib. TUNEL assay to detect apoptotic DNA fragmentation indicated no positive cells (green fluorescence) in OE and KO tumors, indicating no apoptotic cells (Fig.8D, Supplementary Fig. S11). Tumors from olaparib-treated mice exhibited a few TUNEL-positive cells present in OE tumors, whereas the majority of cells were TUNEL-positive in KO tumors. Tumors were assessed for presence of γH2AX in tissue sections by immunohistochemistry (Fig. 8E). Vehicle treated mice exhibited low levels of γH2AX in both KO and OE cohort (Fig. 8E). Tumors from olaparib-treated mice exhibited low levels of γH2AX expression in OE, whereas the majority of cells were positive in KO group (Fig. 8E). These findings indicate a large number of DNA double-strand breaks were generated due to suppression of LCK and inhibition of PARP. We next assayed for expression of CD31, an indicator of microvessel (angiogenesis) density and of tumor mass and growth. Vehicle treated mice exhibited high CD31 positive staining in both KO and OE cohort (Fig. 8F). Tumors from olaparib-treated mice exhibited high levels of CD31 expression in OE, whereas there was no detectable CD31 in the KO group (Fig. 8F). These findings suggested that olaparib was also sufficient to inhibit tumor angiogenesis, corroborating previous findings which showed that PARP facilitates tumor vascularization by augmenting CD31 and VEGF^28^.

## Discussion

Endometrioid ovarian cancer, while rare, lacks effective therapeutic strategies and are HR proficient. Our studies identified a strategy to induce HR deficiency in endometrioid ovarian cancer. We discovered that inhibition or disruption of the non-receptor tyrosine kinase LCK attenuates the expression of HR proteins RAD51, BRCA1, and BRCA2 in eEOC. This complements our previous study showing that LCK overexpression or LCK inhibition modulated the mRNA levels of HR DNA repair genes including RAD51, BRCA1, and BRCA2 ^15^. Here, we showed that LCK modulates HR genes at the protein level. This leads to functional consequences as the inhibition of LCK, via shRNA or pharmacologic inhibitor, leads to inhibition of DNA damage repair as assessed using the established DR-GFP assay in U2OS osteosarcoma cells. LCK does not impact PARP and NHEJ repairs, the alternate mechanisms for repair of double strand breaks by direct ligation independent of an homologous template^21^. The NHEJ repair proteins, Ku70 and Ku80, did not have their expression levels impacted by PP2 treatment. LCK kinase activity is necessary for maintaining HR proficiency. Finally, we demonstrated that LCK disruption is sufficient to sensitize endometrioid ovarian cancer cells to olaparib. These findings are consistent with our data indicating that LCK inhibition leads to chemosensitization to cisplatin treatment in endometrioid ovarian cancer.

We determined for the first time that the LCK protein is upregulated in response to DNA damage in eEOC. DNA damage by etoposide, methyl methanesulfonate (MMS), and by UV radiation induces LCK protein expression. This is also corroborated by previous findings that indicate fractionated radiation induces stem cell populations in human gliomas to display LCK activation ^29^. Phosphorylation of LCK (pY394) was elevated in etoposide, MMS, and UV-treated SKOV3 cells and in UV-treated CP70 cells as compared to untreated cells. We found that DNA damage led to nuclear localization of both total and pY394 LCK protein, a finding supported by immunofluorescence analysis of pY394 LCK following DNA damage. This finding is unprecedented as LCK is localized to the inner leaflet of the cell membrane on microdomains^30^. Previous studies have found constitutively active LCK in the nucleus where it binds at the promoter region of LIM domain only 2 (LMO2) leading to gene expression^31^. Our findings are significant as we show that LCK is activated by DNA damage, leading to nuclear translocation and subsequent activation of HR repair pathways.

Here we confirm nuclear translocation of LCK in response to DNA damage as shown by co-immunoprecipitation with BRCA1 and RAD51. Interaction of LCK with BRCA1 and RAD51 requires kinase activity or phosphorylation on Y394, indicating active LCK is necessary for complex formation^23^. In contrast, phosphorylation on Y192 is not required for complex formation. Moreover, kinase activity and autophosphorylation are necessary for functional DNA repair, as shown by γH2AX and RAD51 foci formation assays. These findings indicate that LCK kinase activity and autophosphorylation is essential to allow for interaction with HR repair proteins BRCA1 and RAD51 during DNA damage response.

LCK regulates HR repair in response to DNA damage and its inhibition potentiates the activity of PARPi to induce synthetic lethality. The simultaneous inhibition of LCK and PARP with pharmacological agents PP2 and olaparib showed significantly more DNA damage and chromosomal aberration compared to only either PP2 or olaparib treatment in eEOC cells. PP2 treatment was sufficient to attenuate DNA repair, augmenting the effect of olaparib in ovarian cancer cells. Finally, in vivo studies showed that olaparib efficacy was enhanced in CP70 LCK KO compared to OE tumor bearing mice. This provides evidence for proof of concept for utility of LCK inhibitors to disrupt HR DNA damage repair. Indeed, several strategies are currently being explored in the clinic to increase use of PARP targeted therapy in HR proficient cancers. CDK1 and CDK12 inhibition led to HR deficiency by decreasing HR repair proteins RAD51, BRCA1, and BRCA2 in lung cancer^8^. Further, inhibition of BET proteins also led to attenuation of RAD51 and BRCA1 proteins in breast, ovarian, and prostate cancer models ^2^. PI3K inhibition is also sufficient to reduce BRCA1 and BRCA2 expression, hampering HR repair in triple-negative breast cancer^32^. Other reported targets are HSP90 ^33^ and VEGFR3 ^34^ to attenuate RAD51, BRCA1, and BRCA2 expressions in ovarian cancer. Concurrent inhibition of LCK enhanced the efficacy of PARPi in above HR proficient cancer models in preclinical settings. Clinical trials are now going on to assess the efficacy of PARPi in combination with CDK1/12 inhibitors, PI3K inhibitor, and VEGFR3 inhibitors^3^. Our findings complement these studies and identify a new signaling pathway for enhancing PARP targeted therapy in eEOC.

These findings provide an innovative new strategy for inducing an HR-deficient status in an otherwise HR-proficient tumor. We identify targeted therapies that compromise HR repair genes and augment sensitivity to PARPi. Our study defines the mechanistic impact of LCK and potentially other non-receptor tyrosine kinases in regulation of HR repair that is apparently crucial to ovarian cancer’s response to chemotherapy and PARP inhibitors. This study highlights new clinical applications that target LCK, expanding PARPi utility.

## Methods

### Cell lines and culture conditions

Cisplatin resistant eEOC cancer cells CP70 were a gift from Analisa Difeo (University of Michigan) and SKOV3 were purchased from American Type Culture Collection (ATCC). Others cell lines used in this study are mentioned in resource table. Cells were grown in DMEM and McCoy’s 5A media respectively, supplemented with 10% fetal bovine serum at 37°C in humidified incubator in 5% CO2. Cells were tested and confirmed as mycoplasma contamination negative on a quarterly basis. Cells were passaged by treatment with trypsin/EDTA solution when they reached 80-90% confluence and further passaged or seeded for experiments.

### Chemicals and reagents

We used a number of pharmacological agents in our study. The PARP inhibitor (Olaparib)^35^, the LCK inhibitor (PP2)^17, 18^ and the radiomimetic drug, etoposide^36^ were purchased as shown in the resource table. Inhibitors were dissolved in 100% DMSO to make stock concentrations and kept at −20°C until use. The details of chemical, reagents, primary antibodies, and secondary antibody details are outlined in the resource table.

### Plasmid construct mutants

Myc-tagged LCK containing plasmid was generated using pENTR/D-TOPO cloning kit (Thermo Scientific) according to manufacturer instructions. Briefly, Myc-LCK gene block was purchased from Integrated DNA Technologies (IDT, USA). Myc-LCK was cloned into pENTR/D-TOPO vector. The entry clone was further transferred into a destination vector, pLenti CMV Puro DEST (Addgene). The plasmid was validated by DNA sequencing (Eurofins). LCK mutants 192F, Y394F, 273R were generated by site directed mutagenesis and sequenced. Each mutant was cloned into a lenti viral vector, pLenti CMV Puro DEST (Addgene) for subsequent use.

### Lentivirus production

Lentiviral particles for LCK silencing were generated using established lab protocols^15^. Briefly HEK293T cells were seeded into 6 well plates. The next day cells were transfected with pRSV-Rev, pMDLg/pRRE, pMD2.G and lentivral vector expressing shRNA for targeting LCK (KD1, TRCN0000426292, KD2, TRCN0000001599). Following 24h incubation, transfection media was replaced with fresh DMEM medium. 48h post transfection, lentiviral particle containing media was filtered to remove cell debris and added to CP70 and SKOV3 cells. Fresh media was subsequently added to the HEK-293T transfection plates and incubated for an additional 24 hours followed by filtration and addition to further CP70 and SKOV3 cells. Transduced CP70 and SKOV3 cells were identified using 1.5ug/ml and 2ug/ml puromycin (Thermo Scientific) selection respectively.

### Generation of CRISPR/Cas9 KO cells

CP70 cells were used to generate LCK CRISPR/Cas9 knockout cells according to the manufacturer protocol (Santa Cruz Biotechnology). Briefly, cells were transfected with GFP labelled LCK CRISPR/Cas9 plasmid using lipofectamine 3000 (Thermo Scientific) in the presence of antibiotic-free, FBS-enriched, Optimem media. Following transfection, cells were kept in transfection medium for 24h, then replaced with fresh culture media. After an additional 24h, transfected cells were screened for GFP expression using a flow cytometer, and the GFP^+/high^ population was isolated and plated as single cells into a 96 well plate. Cells were grown and expanded in accordance with standard culture techniques as stated above, followed by western blotting for LCK protein expression with anti-LCK antibody (0.5 µg/mL, R & D Systems). Clones with the lowest LCK expression compared to parental cells were considered LCK KO cells.

### Western blot analysis

Western blot analysis was performed as reported with modifications as follows^37, 38^. Briefly, cancer cells were washed with chilled Dulbecco’s phosphate buffered saline (PBS) two times at the end of treatment. NP-40 lysis buffer (Invitrogen) was added dropwise to the plates and placed on ice for 10 minutes. The NP-40 lysis buffer contains 50 mM Tris, pH 7.4, 250 mM NaCl, 5 mM EDTA, 50 mM NaF, 1 mM Na3VO4, 1% Nonidet™ P40 (NP40), 0.02% NaN3 and was supplemented with 1mM PMSF and 2 µg/ml protease cocktail inhibitor (PCI) (Sigma Aldrich). Cells were then collected in a 1.5 mL centrifuge tube by scraping, and incubated on ice for one hour with occasional vortexing. Lysates were centrifuged at 10,000rpm for 10 min at 4°C. Protein concentration was measured of each lysate supernatant by BCA kit analysis (Thermo Scientific). Protein samples were then prepared using 6x Laemmli dye containing BME (β-mercapto ethanol) and boiled for 5-10min. Protein samples were subjected to SDS-β PAGE gel electrophoresis using pre-made gradient gels (4-20%, Biorad). Proteins were transferred by wet transfer to a PVDF membrane (Millipore) at 4°C overnight. Membranes were then blocked in 5% BSA in TBST for one hour at room temperature, and subsequently, incubated overnight at 4°C in the following primary antibodies: T-LCK (1:1000 R&D Systems), T-LCK (1:1000 Proteintech), P-LCK 394 (1:1000 R&D Systems), RAD51 (1:1000, Proteintech), BRCA1 (1:500, EMD Millipore), BRCA2 (1:500, EMD Millipore), γH2AX (1:1000 Cell Signaling Technology), GAPDH (1:5000 Proteintech), and β-actin (1:4000 Proteintech). After primary antibody incubation membranes were washed three times with TBST (Tris-buffered saline containing 0.1% tween 20) washing buffer on a platform shaker. Membranes were incubated with HRP-conjugated rabbit (1:25000) or mouse (1:25000) secondary antibodies for one hour at room temperature, followed by three washes with TBST buffer. Chemiluminescence reagent (PerkinElmer) was added to detect immobilized proteins in PVDF membranes utilizing the ChemiDoc imaging system. Densitometry was performed using Image J software.

### Nuclear protein isolation and co-immunoprecipitation analysis

CP70 and SKOV3 cells transduced with Myc-tagged LCK were treated with etoposide (10µM) or DMSO for 24h followed by replacement with fresh serum-enriched media for an additional 24h. Cells were collected and washed with cold PBS two times, scraped and centrifuged. Cell pellets were then lysed with cytoplasmic and nuclear extraction buffers according to manufacturer protocols (NE-PER Nuclear and Cytoplasmic Extraction Kit, Thermo Scientific). Protein concentrations of nuclear lysates were estimated using the BCA method outlined above. For co-immunoprecipitation, nuclear protein lysates were incubated with 3µg anti-Myc antibody (Proteintech) or 3µg control antibody (Cell Signaling Technology) overnight at 4°C with gentle rocking. Pre-cleaned protein A/G agarose beads (Thermo Scientific) were added to the lysates and incubated for 4h at 4°C on a rotating mixer. Beads were then collected by centrifugation and washed three times with chilled NP-40 lysis buffer. 6x Laemmli buffer (Alfa Aesar) containing BME was added and beads were boiled for 5 minutes. Samples were separated on SDS-PAGE and processed for western blot analysis as outlined above. Further, LCK overexpression (OE) SKOV3 (without Myc tagged) cells were treated with etoposide (10µM) for 24h. Then, serum-enriched media was added to replace drug-containing media and kept for another 24h. Then, cells were collected, and nuclear lysates were prepared. Further immunoprecipitation/co-immunoprecipitation was performed after RAD51 pulled down as described above.

### Gene conversion assay

Gene conversion assay or DR-GFP assay was performed according to reported methods^39^. Human osteosarcoma U2OS cells stably transfected with DR-GFP plasmid and I-SceI endonuclease expression vector pCBASce were kindly provided by Maria Jasin at Memorial Sloan-Kettering Cancer Center. Cells were treated with, PP2 (5, 7, and 10µM) or DMSO vehicle for 48h. Cells were then transfected with I-SceI endonuclease expression vector pCBASce using Lipofectamine 3000. In a separate set of experiments, U2OS cells with DR-GFP integration were transfected with shCon, LCK KD1 or KD2 for 24h and incubated in serum enriched medium for another 24h. Cells were further transfected with I-SceI plasmid for 24h followed by incubation with serum enriched medium for 24h. Live cells (Live/Dead dye kit, Thermo Scientific) were analyzed with a flow cytometer to estimate the percentage of GFP-positive cells.

### RAD51 and **γ**H2AX nuclei staining and analysis

Laser scanning confocal microscopy was performed to detect RAD51 and γH2AX foci in cancer cells. Briefly, ovarian cancer cell, CP70 were grown on coverslips and treated with 10µM etoposide or vehicle for 24h followed by an additional 24h in drug-free media. Coverslips were washed with PBS and fixed with 4% paraformaldehyde (Electron Microscopy Sciences) in PBS. Cells were permeabilized with 0.01% triton-X 100 (Fisher Scientific) for 5 minutes followed by a wash with chilled PBS and blocked with 3% goat serum (Thermo Scientific) for 1h at room temperature. Cells were incubated with anti-RAD51 (1:250, Abcam) or anti-γH2AX (1:300, Cell Signaling Technology) antibodies overnight at 4°C in a humidified chamber. Next, cover slips were washed 3X with PBS. Alexa fluorescent conjugated secondary antibodies were added to the coverslips and incubated for 1h. Coverslips were washed 3x and mounted with DAPI containing Vectashield (Vector Lab). Images were captured by confocal microscope at 63x magnification in oil emersion (Leica SP8 confocal microscope). RAD51 and γH2AX foci were counted on 20 representative cells by image J software.

### Metaphase spread analysis

Metaphase spread analysis was performed on LCKi and PARPi treated cells using established methods ^27^. Briefly, cells were treated with LCK inhibitor, PP2 (5µM) (Selleck Chemicals) and PARP inhibitor, olaparib (3µM) (Selleck Chemicals) for 48h. After treatment, cells were harvested. Cells were treated with colcemid (50ng/ml) (Sigma) for 1.5h then washed with PBS and placed in 0.075 mol/L KCl (Sigma) solution for 20min. Subsequently, cells were washed with PBS and fixed in carnoy fixative solution (Methanol: acetic acid 3:1) added dropwise followed by one hour incubation. Cell pellets were collected, and fixative solution was added and incubated at 4°C for 24h. Cell pellets were collected and small amount fixative solution was added and cell suspension was slowly dropped on glass slides and allowed to dry at 37°C. Slides were then stained with Giemsa solution (Sigma Aldrich). Images were captured at 100X magnification with a bright field microscope. Abnormalities in chromosomes were quantified (Chromosomal break, gap, radial formation) by visually counting five nuclei per treatment group.

### Single Cell Electrophoresis Assay

Single cell electrophoresis assay or comet assay was performed according to a previously reported method^40^. This experiment was performed following the manufacturer’s instructions (Trevigen). Briefly, CP70 and SKOV3 eEOC cells were treated with 5µM PP2 and/or 3µM Olaparib for 48h. Comet LMAgarose was melted at 90°C for 10min in a water bath then cooled for 20min to 37°C. Treated cells were detached from plates using trypsin. Serum enriched media was added to neutralize the trypsin. Cell suspension was washed twice with chilled 1X Ca^++^ and Mg^++^ free PBS, and subsequently suspended at 1 × 10^5^ cells/ml in chilled 1X PBS buffer (Free of Ca++ and Mg++). For the alkaline comet assay, the cell suspension was mixed with molten LMAgarose at 37°C. Immediately, 50µL LMAgarose mix was spread on glass microscope slides and incubated at 4°C for 30 min in the dark. Slides were incubated in lysis solution (provided in kit) overnight at 4°C. The next day comet slides were incubated in alkaline unwinding solution at room temperature for 20 min. Agarose gel electrophoresis (21volts for 30min) was performed using alkaline electrophoresis protocol. After electrophoresis, slides were briefly immersed in distilled water twice and then immersed in 70% ethyl alcohol for 5 min. Slides were dried and 100µl of SYBR gold (excitation/emission is 496 nm/522 nm) was added on agarose and kept for 30 min in the dark. Slides were washed with water, dried, and visualized by fluorescence microscopy.

### Colony formation assay

The pharmacological effect of olaparib in cancer cells was investigated by colony formation assay according to the earlier reported method^27^. Briefly, ovarian cancer cells CP70 and SKOV3 cells (WT, sh, LCK OE, LCK KO, LCK KD and mutants) were placed on 12 well plates. The next day, cells were treated with olaparib in a dose dependent manner for 12 days. During this time the media was changed every day using fresh drug. At the end of experiment, PBS was added to the well to wash the colony. Then, cells were fixed with 4% paraformaldehyde solution for 10min. Cells were then washed two times with PBS and incubated in 0.2% crystal violet solution for one hour at room temperature. After incubation cells were washed three times with PBS. Then, the images of six well plates were captured and number of colony forming area was analyzed by Image J software.

### Cell titer glo viability assay

CP70 (WT, OE, KD, KO and LCK mutants, Y394F, K273R and Y192F) cells were collected after trypsinization. Cells were counted and 4000 cells were plated in each well of a 96 well plate. Cells were then treated with etoposide for 48h in a dose dependent manner. Control cells were treated with vehicle (DMSO). After the drug treatment, Cell TiterGlo® Luminescent Cell viability assay reagent (Promega) mixture was prepared and added to the cells to lyse them for 10 minutes shaking and luminescence was measured via luminometer. Cell viability percentage was calculated as luminescence of treated group/luminescence of vehicle treated group×100.

### NHEJ gene expression

To check for NHEJ expression, CP70 and SKOV3 cells were grown on 60mm petri dishes until 70% confluent. Cells were then treated with PP2 in a dose dependent manner (5, 7 and 10µM) for 48h and collected by scraping and Western blot was performed to assess protein expression of NHEJ markers Ku70 and Ku80.

### *In vivo* animal study in NSG mice

*In vivo* antitumor efficacy of Olaparib was tested in NSG (NOD severe combined immunodeficient (SCID) IL2R gamma) mice. This study was approved (IACUC Protocol# 2707) by Institutional Animal Care and Use Committee (IACUC), Cleveland Clinic Lerner Research Institute. Animals were procured from BRU (Biological Response Unit) facility of the Cleveland Clinic. All cells used in this study were transfected (Lentiviral transfection) with pCDH-EF1a-eFFly-mCherry plasmid. Mice were injected with CP70 LCK knockout cells or CP70 LCK overexpression cells intraperitoneally (Cells: 0.5×10^6^). After that mice were divided into four groups-**1**. Mice: CP70 LCK KO: Vehicle (n=8), **2**. Mice: CP70 LCK KO: Olaparib (50mg/kg) (n=8), **3**. Mice: CP70 LCK OE: Vehicle (n=8), **4.** Mice: CP70 LCK KO: Olaparib (50mg/kg) (n=8). Olaparib was dissolved in ddH_2_O containing 4% DMSO and 30% polyethelene glycol (PEG300) and injected intraperitoneally. Animals were treated with olaparib for 5days/week. Bioluminescence of the tumor were measured by *in vivo* imaging system (IVIS Spectrum CT, PerkinElmer). At the end of the experiment, mice were sacrificed according to the protocol of Institutional Animal Care and Use Committee (IACUC), Cleveland Clinic Lerner Research Institute. Tumor tissues were collected and preserved in 10% formalin solution.

### Immunohistochemistry and TUNEL assay

Immunohistochemical (IHC) analysis and TUNEL assay of tumor sections were performed according to the earlier reported method^41^. Formalin fixed tissues were sent to histology core to make thin slice (around 5micron) of tissue embedded on glass slides. Next, slides were dipped in Histo-Clear to deparaffinize. Then sections were rehydrated in gradient ethanol (100%, 95%, 80% and 60% ethanol 5 min for each bath). For antigen retrieval, sections were then put in Tris-EDTA buffer (pH9) and boiled in a pressure cooker for 10min. Then slides were put for an hour to cool. Slides were then put into 0.1% triton X-100 solution for 10 min to permeabilize. After washing with PBS three times, tissue sections were blocked with blocking buffer (5% goat serum and 0.1% triton X-100 in 1X TBST) at room temperature for one hour. Then antibodies were added to the tissues and incubated for overnight at 4°C. Next day, sections were washed three times with PBS. Then diluted 100 µl Peroxidase Labeled Polymer was added to the tissues and incubated for 30 min at room temperature. Now sections were washed three times with PBS. Then diluted DAB chromogen was added to the tissue and incubated for 2-5min until desired color was generated. Sections were washed three times and then hematoxylin staining was performed. After washing with PBS, tissue sections were dried and mounted with Cytoseal. Images were captured using upright microscope at 20x magnification.

For TUNEL assay, tissue sections were processed for antigen retrieval and permeabilization as discussed earlier. Sections were washed three times with PBS. Then enzyme solution was prepared according to the manufacturer instructions. Slides were then incubated in the enzyme for 60min at 37°C. Sections were washed three times with PBS and mounted with Vectashield containing DAPI.

### Software and Statistics

Graph pad prism software was utilized for graph preparation and to determine statistical significance (detailed in each figure legend). Image J was used for quantification of data. Each experiment was performed at least three times. p-value less than 0.05 was considered significant.

## Acknowledgments

Authors thank the members of Reizes and Lathia Laboratories for helping to improve the scientific quality of the manuscript. We would like to thank Alex Myers for extensive editing the manuscript. We thank Dr. Gauravi Deshpande for helping in image acquisition by confocal microscopy, Amy Graham and Eric Schultz for flow cytometry, and Drs. Debjit Khan and Krishnendu Khan from Fox Laboratory for insights on complex formation assays. We would also like to thank Dr. Suparna Mazumdar (Cleveland Clinic) for giving constructive comments of this study. Research in the Reizes laboratory is funded by Center of Research Excellence in Gynecologic Cancer from the Cleveland Clinic Foundation, VeloSano Bike to Cure, and the Laura J. Fogarty Endowed Chair for Uterine Cancer Research. Dr. Gong is supported by and NIH/NCI grant (R01 CA222195).

## KEY RESOURCES TABLE

| REAGENT or RESOURCE | SOURCE | IDENTIFIER |
| --- | --- | --- |
| **Antibodies** |  |  |
| Anti-LCK antibody | Proteintech | 12477-1-AP |
| Anti-LCK antibody | R&D System | MAB3704 |
| Anti-P-LCK (Y394) antibody | R & D system | MAB7500 |
| Anti-RAD51 antibody | Proteintech | 14961-1-AP |
| Anti-RAD51 antibody | Santa Cruz | sc-398587 |
| Anti-P-H2A.X (Ser139) Antibody | Cell signaling | 2577S |
| Anti-BRCA1 antibody | EMD Millipore | OP-92-100UG |
| Anti-BRCA2 antibody | EMD Millipore | OP-95-100UG |
| Anti-Myc antibody | Proteintech | 60003-2-Ig |
| Anti-GAPDH antibody | Proteintech | HRP-60004 |
| Anti-Lamin A/C antibody | Proteintech | 10298-1-AP |
| Alpha Tubulin Monoclonal antibody | Proteintech | HRP-66031 |
| Anti-Ku70 antibody | Proteintech | 10723-1-AP |
| Anti-Ku80 antibody | Proteintech | 16389-1-AP |
| Rabbit IgG XP® Isotype Control | Cell signaling | 3900S |
| Mouse IgG XP® Isotype Control | Cell signaling | 5415S |
| Anti-Rabbit IgG (H+L), HRP antibody | Promega | W4018 |
| Anti-Mouse IgG (H+L), HRP antibody | Promega | W4028 |
| Goat anti-Rabbit IgG Alexa Fluor 488 | Thermo | A32731 |
| Goat anti-mouse IgG Alexa Fluor 568 | Thermo | A11031 |
| **Chemicals and reagents** |  |  |
| Precision Plus Protein™ Kaleidoscope | Biorad | 1610375 |
| Olaparib | Selleck chemicals | S1060 |
| PP2 | Selleck chemicals | S7008 |
| Etoposide | Selleck chemicals | S1225 |
| Cisplatin | Fesenius Kabi | 401572I |
| Colcemid | Sigma | 10295892001 |
| Laemmli SDS sample buffer, reducing (6X) | Alfa Aesar | J61337 |
| Pierce™ Protein A/G Plus Agarose | Thermo | 20423 |
| Precision Plus Protein™ Kaleidoscope™ | 1610375 | Biorad |
| SYBR™ Gold Nucleic Acid Gel Stain | Thermo | S11494 |
| Trypan Blue | Fisher scientific | 25900CI |
| Immobilon-P PVDF Membrane | Merck Millipore | IPVH00010 |
| **Critical Commercial Assays** |  |  |
| VECTASHIELD® Mounting Medium | Vector lab | H-1200 |
| NP40 Lysis buffer | Thermo | FNN0021 |
| Pierce™ IP Lysis Buffer | Thermo | 87788 |
| Comet Assay Kit | Trevigen | 4250-050-K |
| NuPAGE™ Protein Gel | Thermo | NP0329BOX |
| 4–20% Mini-PROTEAN Protein Gels | Biorad | 4568096 |
| Pierce BCA Protein Assay Kit | Thermo | 23225 |
| Lipofectamin 3000 | Thermo | L3000001 |
| CellTiter-Glo® 2.0 Cell Viability Assay | Promega | G9241 |
| Live/Dead assay kit | Thermo | L23105 |
| KaryoMAX™ Giemsa Stain Solution | Thermo | 10092013 |
| Giemsa Stain, Modified Solution | 32884-1L | Sigma |
| NP40 lysis buffer | Invitrogen | FNN0021 |
| Protease Inhibitor Cocktail | Sigma | 04693159001 |
| Cytoplasmic and nuclear protein isolation kit | Thermo | 78835 |
| **Experimental Models: Cell Lines** |  |  |
| CP70 | Dr. Analisa Difeo | NA |
| SKOV3 | ATCC | NA |
| U2OS | Dr. Zihua Gong | NA |
| U2OS DRGFP reporter cells | Dr. Zihua Gong | NA |
| HEK293T | ATCC | NA |
| CRL1978 | ATCC | NA |
| **Recombinant DNA** |  |  |
| pLenti CMV Puro DEST | Addgene | NA |
| Myc-LCK pLenti CMV Puro DEST | In house |  |
| Myc-LCK Y394F pLenti CMV Puro DEST | In house | NA |
| Myc-LCK K273R pLenti CMV Puro DEST | In house | NA |
| Myc-LCK Y192F pLenti CMV Puro DEST | In house | NA |
| I-SceI plasmid | Dr. Zihua Gong | NA |
| ShRNA targeting LCK | Sigma Aldrich | TRCN0000426292 |
| ShRNA targeting LCK | Sigma Aldrich | TRCN0000001600 |
| ShRNA targeting LCK | Sigma Aldrich | TRCN0000001598 |
| ShRNA targeting LCK | Sigma Aldrich | TRCN0000001599 |
| LCK CRISPR/Cas9 KO Plasmid | Santa Cruz | SC-400434-KO-2 |
| **Software** |  |  |
| FlowJo | BD Bioscience | NA |
| Graph Pad prism | www.graphpad.com | NA |
| ImageJ | imagej.nih.gov | NA |

## Supplementary File

**Supplementary Fig. S1:**
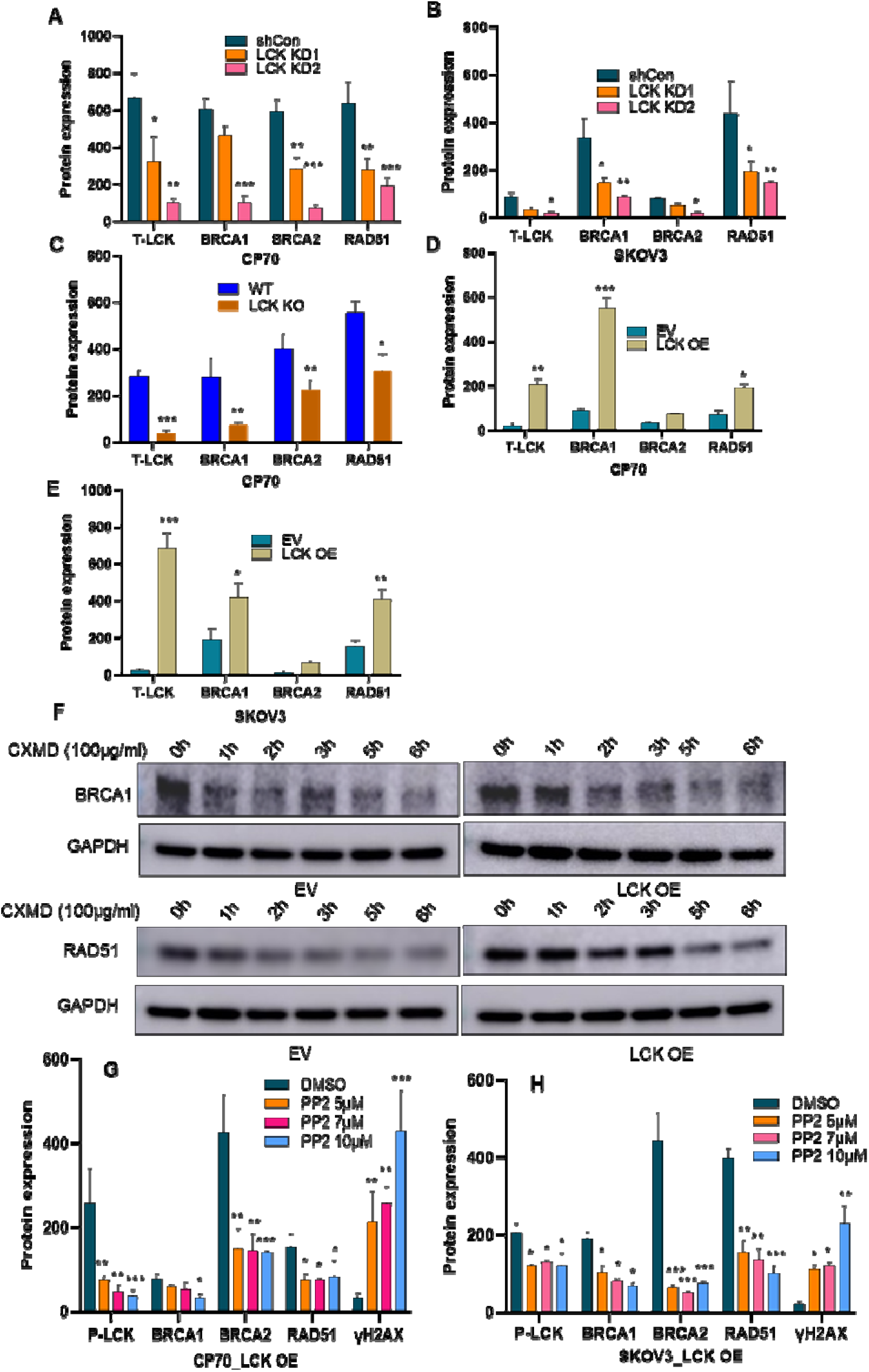
**(A)** Protein quantification T-LCK, BRCA1, BRCA2 and RAD51 in shCon, LCK KD1, and LCK KD2 CP70 cells as observed in western blot analysis (Main Fig.1A, Left panel). **(B)** Quantification of T-LCK, BRCA1, BRCA2 and RAD51proteins in shCon, LCK KD1, and LCK KD2 SKOV3 cells (Main Fig.1A, second panel from left). **(C)** Quantification of T-LCK, BRCA1, BRCA2, RAD51 protein expression in CP70 WT and LCK KO cells (Main Fig.1A, third panel from left). **(D and E)** Quantification of T-LCK, BRCA1, BRCA2 and RAD51 proteins in CP70 EV and LCK OE (Main Fig.1A, fourth panel from left), and SKOV3 EV and LCK OE cells (Main Fig.1A, fifth panel from left). **(F)** Stability study of BRCA1 and RAD51 proteins in CP70 EV and LCK OE cells. **(G and H)** Protein quantification of P-LCK, BRCA1, BRCA2, RAD51 and γH2AX in CP70 LCK OE and SKOV3 LCK OE cells treated with PP2 for 48h in a dose dependent manner (Main Fig. 1D).

**Supplementary Fig. S2:**
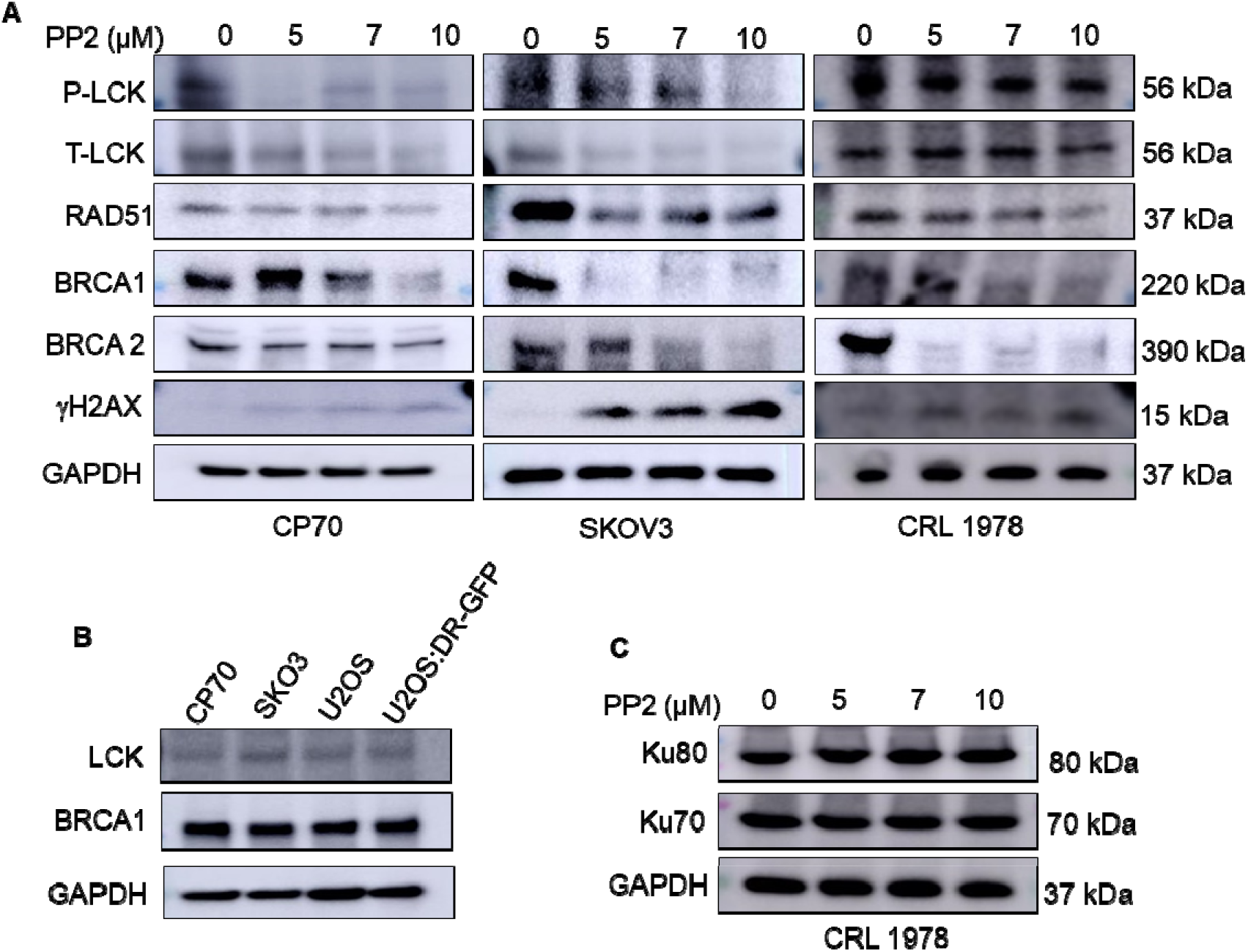
**(A)** Pharmacological inhibition of LCK attenuates HR repair proteins in ovarian cancer cells. CP70, SKOV3 and CRL1978 cells were treated with the LCKi, PP2 for 48h. Cells were harvested, lysed, and analyzed by immunoblot to assess protein expression of BRCA1, BRCA2, RAD51, and γH2AX. **(B)** Western blot analysis in different cells to check the expression of LCK and BRCA1. U2OS is osteosarcoma cell line which was used in DR GFP assay. U2OS and U2OS: DR-GFP cells were examined for checking LCK and BRCA1 expression. These cells were also found to express LCK and BRCA1 like CP70 and SKOV3 cells. **(C)** CRL1978 cells were treated with increasing concentrations of PP2 for 48h and cells were harvested, lysed, and immunoblotted for Ku70, and Ku80 protein expression. GAPDH was used as loading control.

**Supplementary Fig. S3.**
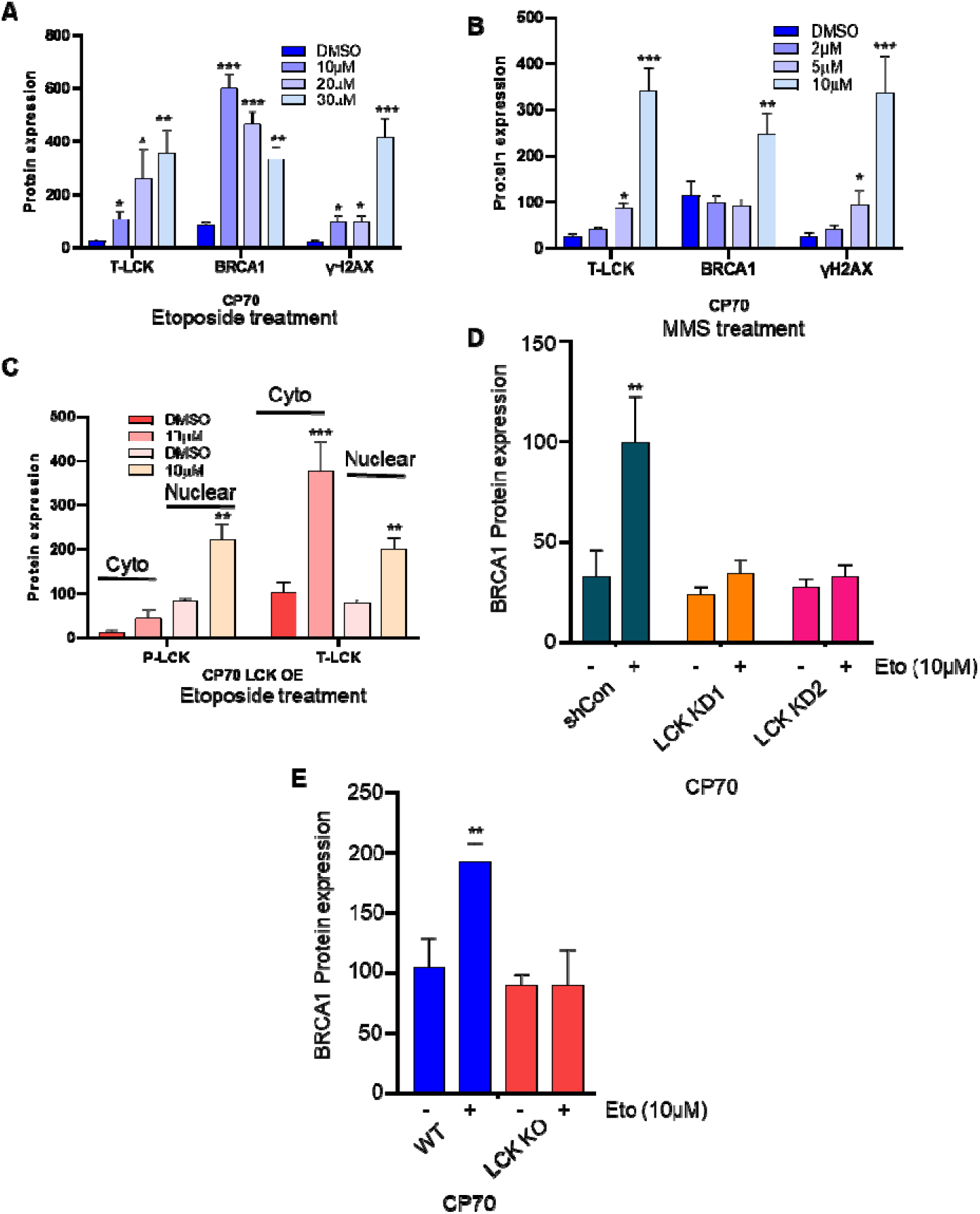
**(A, B)** Protein quantification of T-LCK, BRCA1, and γH2AX in CP70 cells treated with etoposide and MMS (Main Fig. 2A). **(C)** Quantification of P-LCK and T-LCK in CP70 LCK OE cells treated with etoposide (Main Fig. 2B). **(D)** Quantification of BRCA1 protein in CP70 shCon, LCK KD1 and LCK KD2 cells treated with/without etoposide (Main Fig. 3D). **(E)** Quantification of BRCA1 protein expression in CP70 WT and LCK KO cells treated with/without etoposide (Main Fig. 3E).

**Supplementary Fig. S4:**
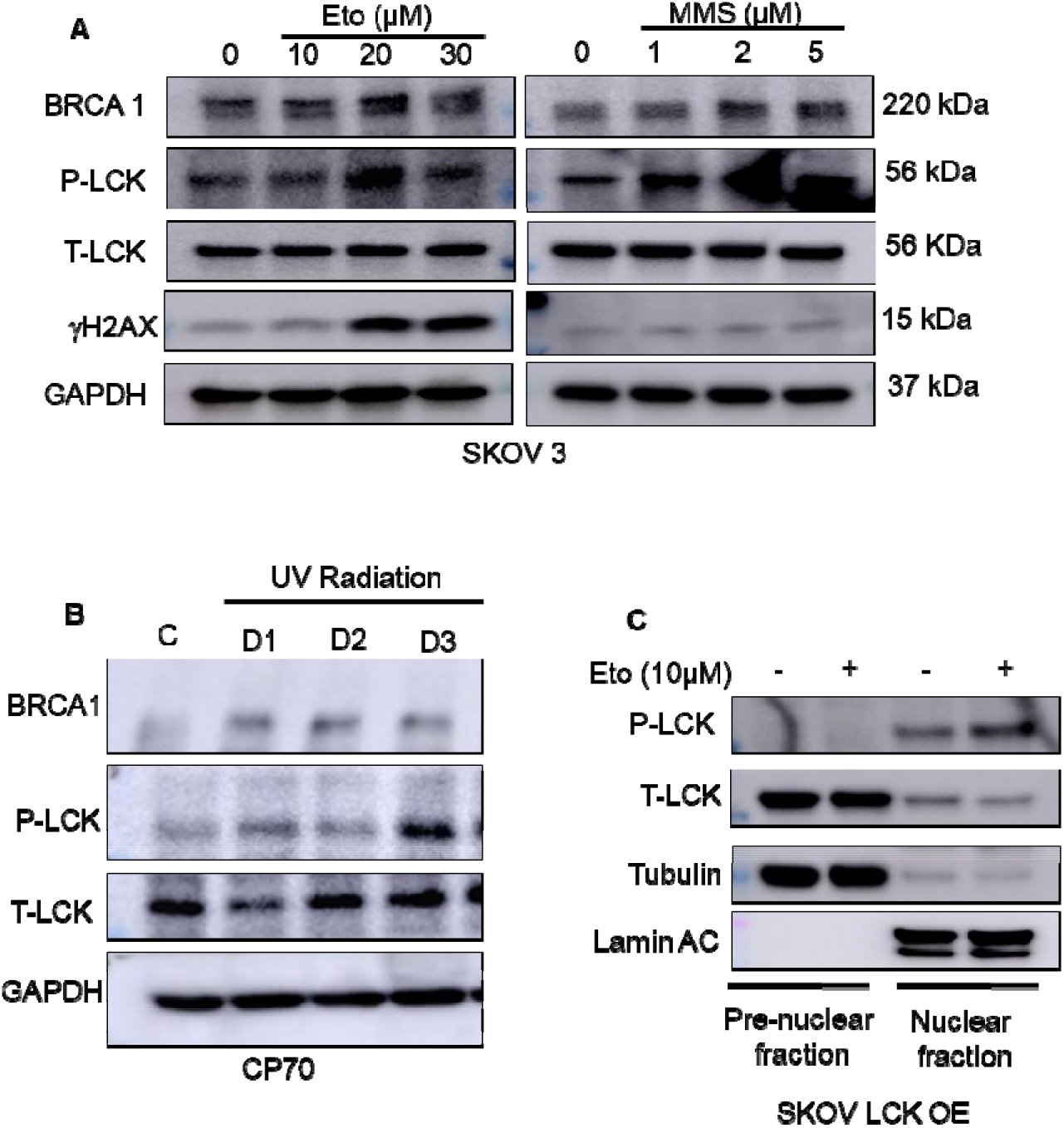
**(A)** SKOV3 cells were treated with etoposide, and MMS for 24. After that cells were put in 24h in drug free media. Cells were then subjected to western blot analysis for checking protein expression. **(B)** DNA damage by UV radiation upregulates LCK phosphorylation. CP70 cells were treated with UV radiation for 1min, 2min and 4min. Cells were kept in serum enriched media for 24h. Then cells were subjected to western blot analysis to check the expression of P-LCK, T-LCK and BRCA1 expression. **(C)** SKOV3 LCK OE cells were treated with etoposide for 24h. Cells were then put in drug free media for another 24h. Cells were collected, and cytoplasmic and nuclear proteins were extracted for western blot analysis.

**Supplementary Fig. S5.**
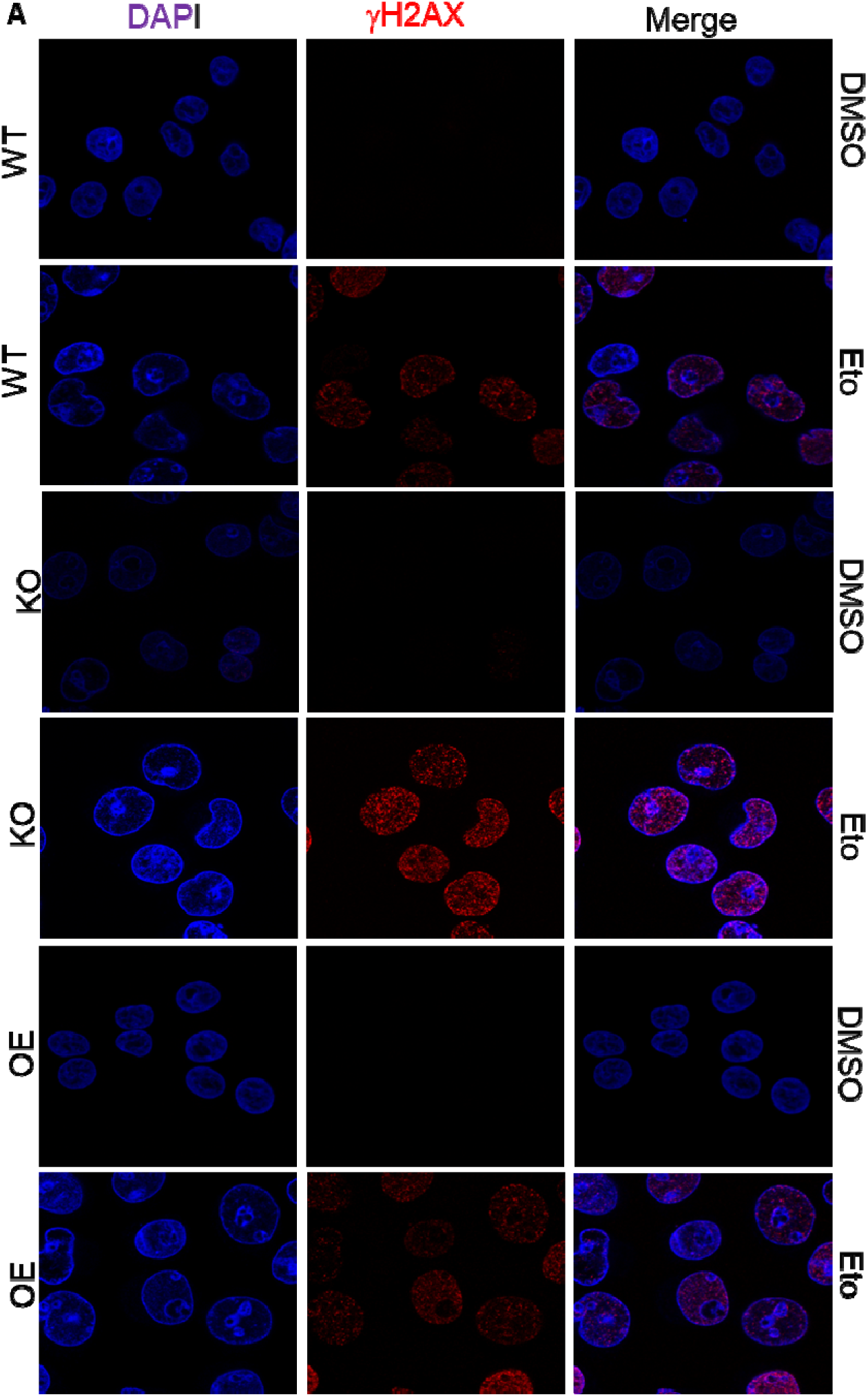
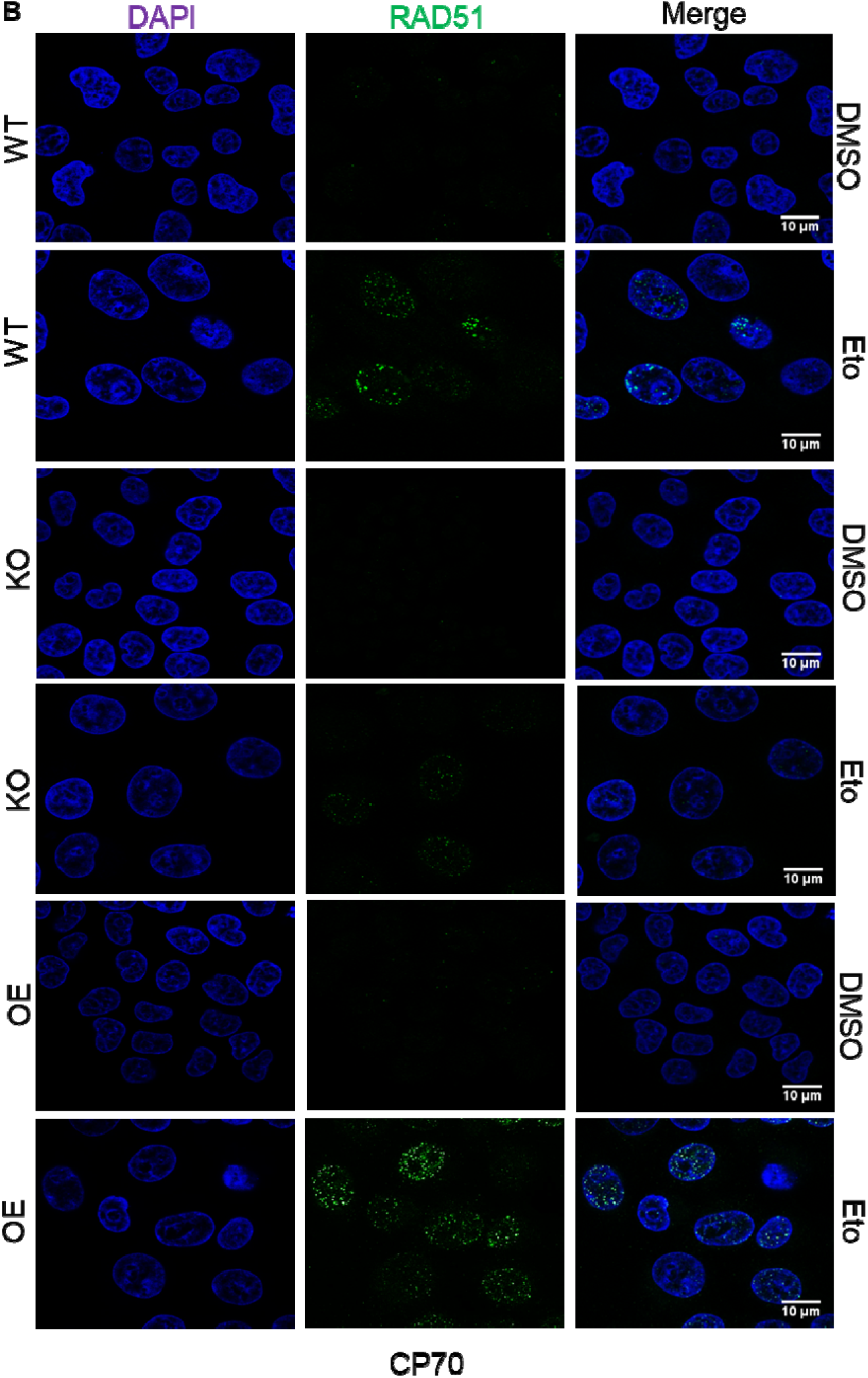
**(A)** CP70 WT, LCK KO (CRISPR/Cas9) and LCK OE (In CRISPR background) cells were treated with DMSO/etoposide 10µM for 24h. Then cells were kept in drug free media for another 24h. Then immunofluorescence study was performed to visualize γH2AX foci formation in different groups. **(B)** CP70 WT, LCK KO (CRISPR/Cas9) and LCK OE (In CRISPR background) cells were treated with DMSO/etoposide 10µM for 24h. Cells were put in drug free media for another 24h. Then immunofluorescence study was performed to visualize RAD51 foci formation in different groups.

**Supplementary Fig. S6:**
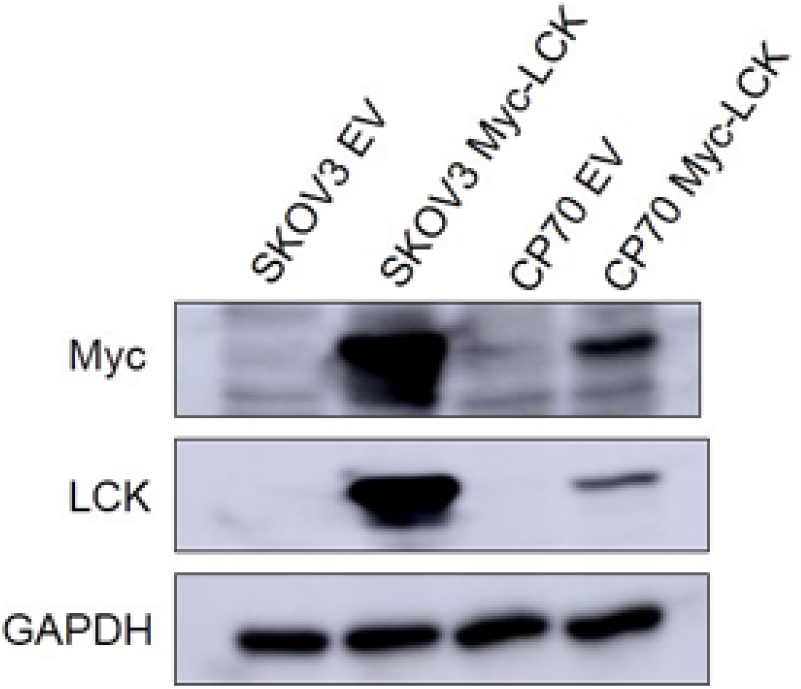
Transfection efficiency of Myc tagged LCK in SKOV3 and CP70. SKOV3 and CP70 cells were transduced with EV or Myc tagged LCK plasmid by using lentiviral particle. Then, cells were checked for Myc and LCK expression.

**Supplementary Fig. S7:**
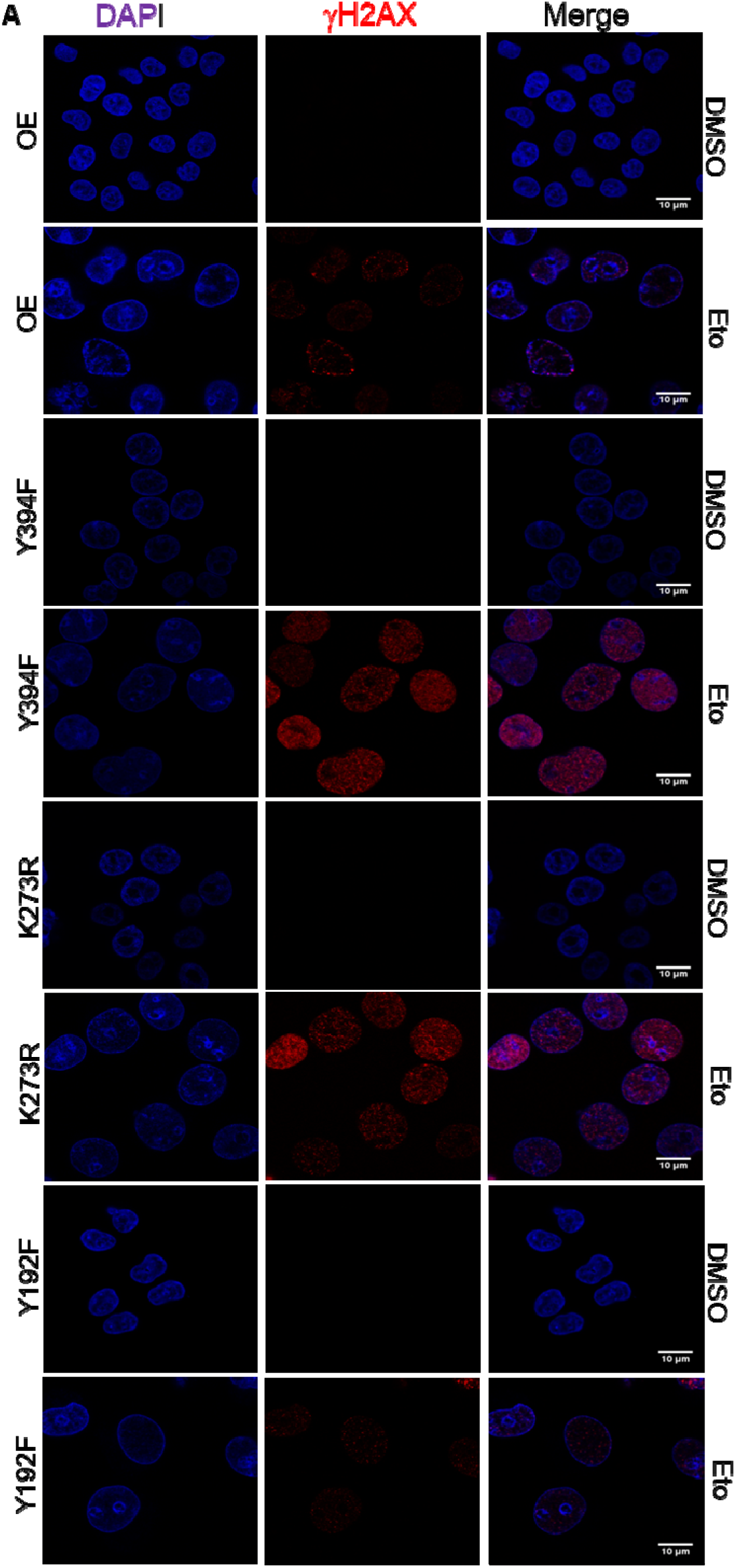
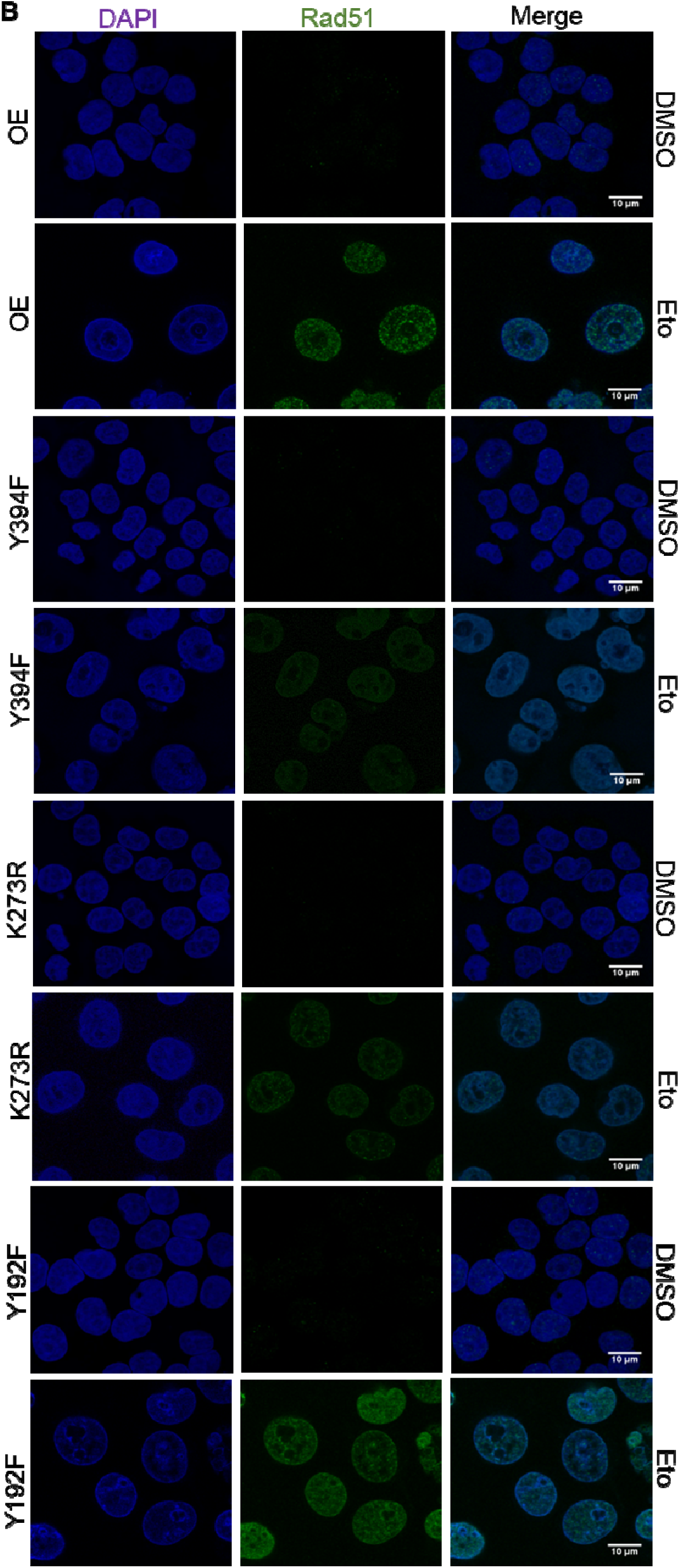
**(A)** CP70 cells (LCK OE, LCK Y394F, LCK K273R, and LCK Y192F, all constructs were introduced into the CP70 LCK KO cells) were treated with etoposide for 24h. Cells were then kept in drug free media for 24h. Immunofluorescence study was performed to visualize γH2AX foci formation. **(B)** CP70 cells (LCK OE, LCK Y394F, LCK K273R, and LCK Y192F) were treated with etoposide for 24h. Cells were then kept in drug free media for 24h. Immunofluorescence study was performed to visualize RAD51 foci formation.

**Supplementary Fig. S8:**
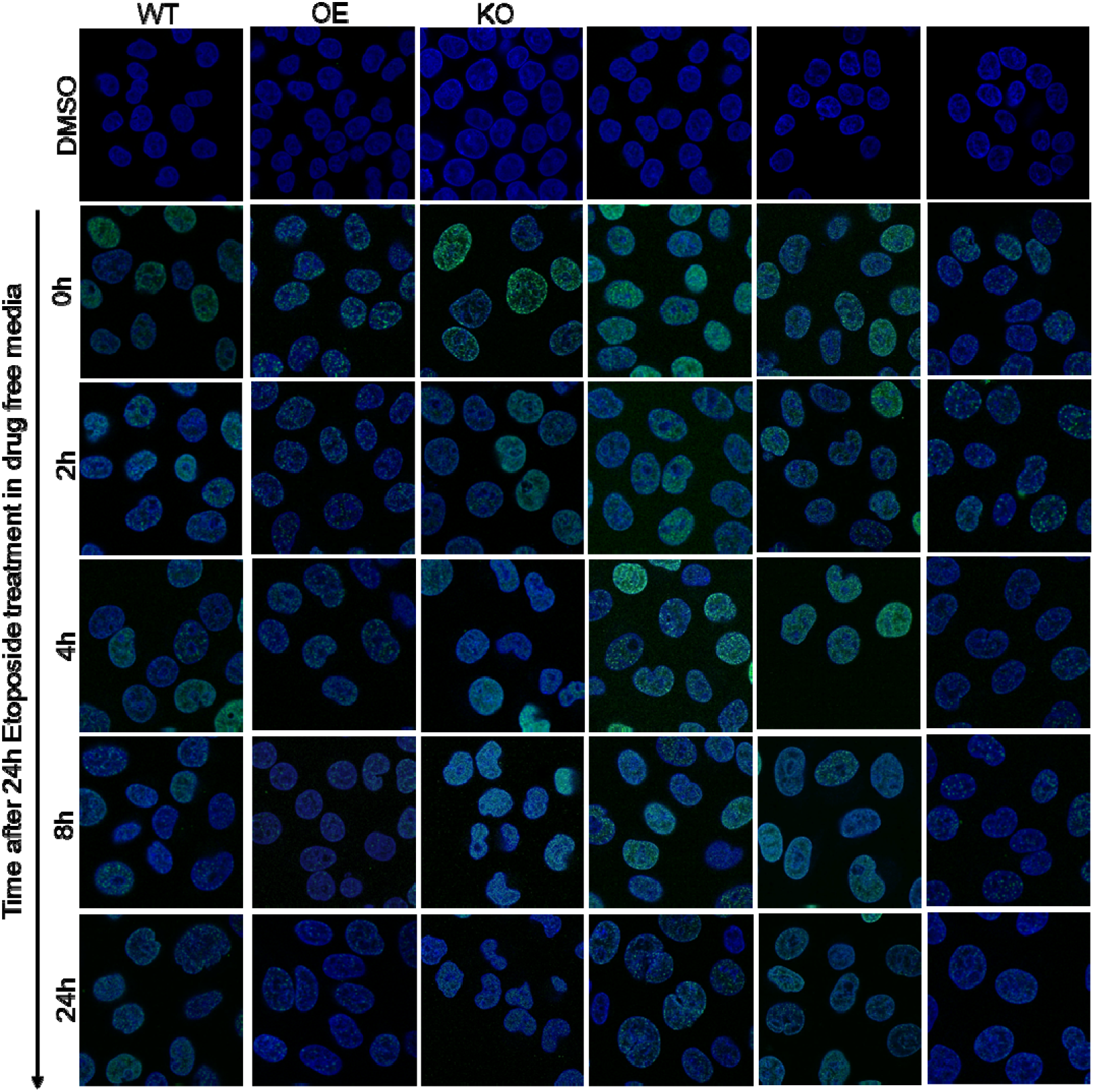
CP70 cells (LCK, LCK Y394F, LCK K273R, and LCK Y192F in LCK knock out background) were grown on cover slips and treated with etoposide for 24h followed by incubation for 0, 2, 4, 8 and 24h. Cells were then subjected to immunofluorescence analysis to visualize H2AX foci formation.

**Supplementary Fig. S9:**
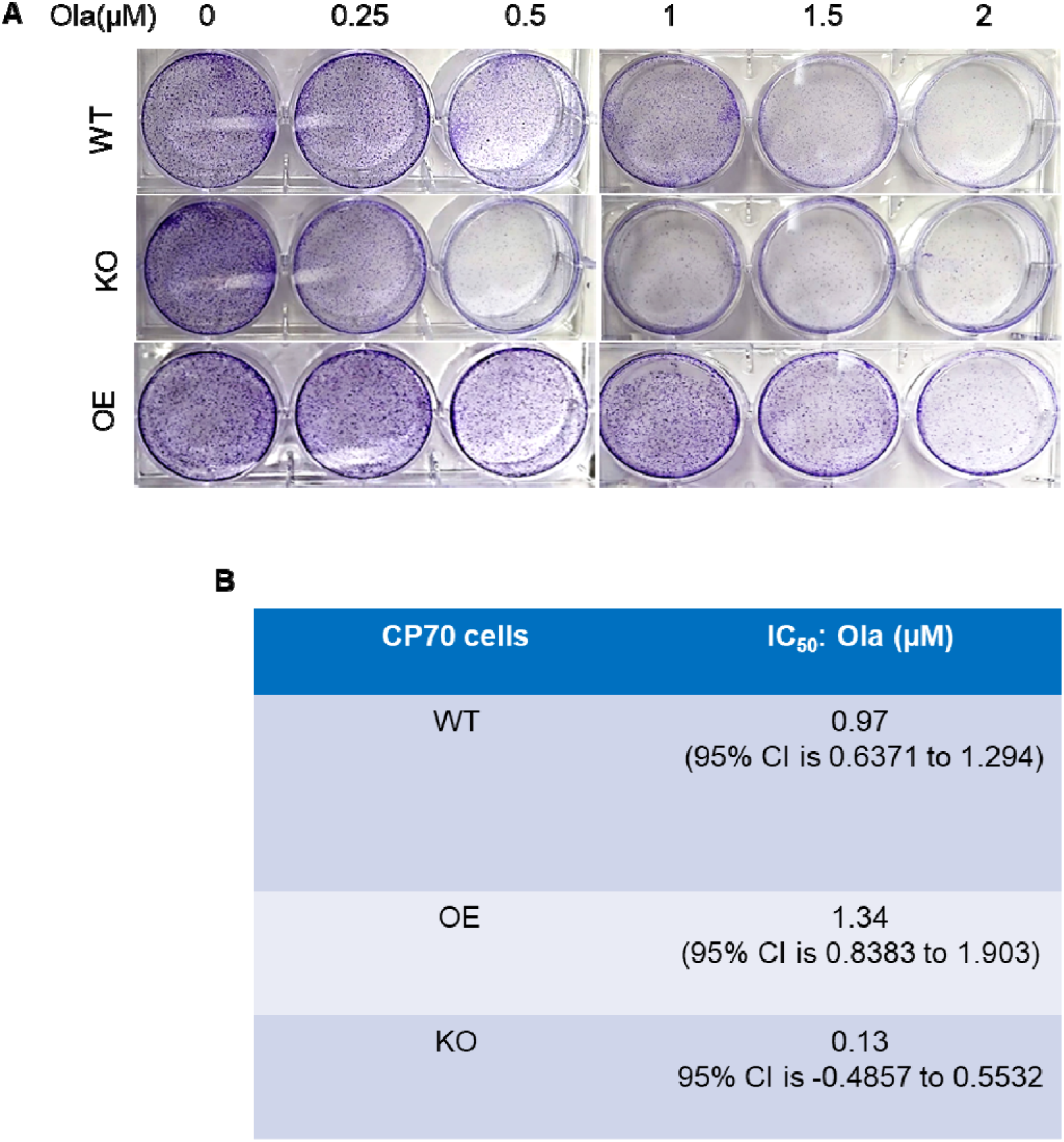
**(A)** CP70 Parental cells and CP70 LCK KO and CP70 CD55 OE (In KO background) cells were treated with Olaparib in dose dependent manner for 12 days. After that colonies were stained with crystal violet and images were captured. **(B)** Number of Colony formation was counted and plotted as percentage of colony formation in the graph (Main fig 6G). IC50 values were shown in the table.

**Supplementary Fig. S10:**
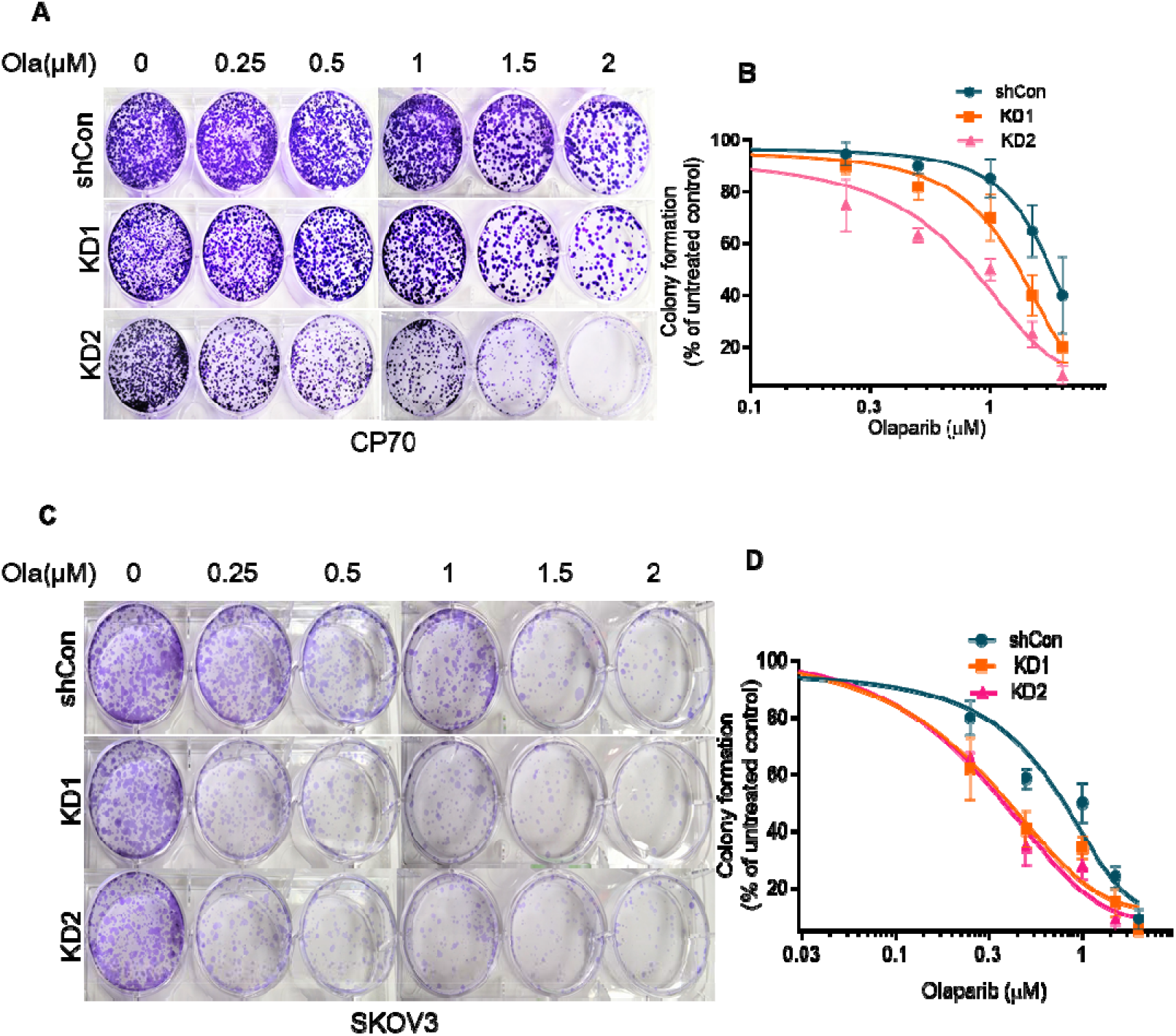
**(A)** CP70 Sh Con or LCK knock down cells were treated with Olaparib in a dose dependent manner for 12 days. Number of colonies was counted and plotted using graph pad prism. **(B)** SKOV3 Sh Con or LCK knock down cells were treated with Olaparib in dose dependent manner for 12 days. Number of Colony formation was counted and plotted in the graph.

**Supplementary Fig. S11:**
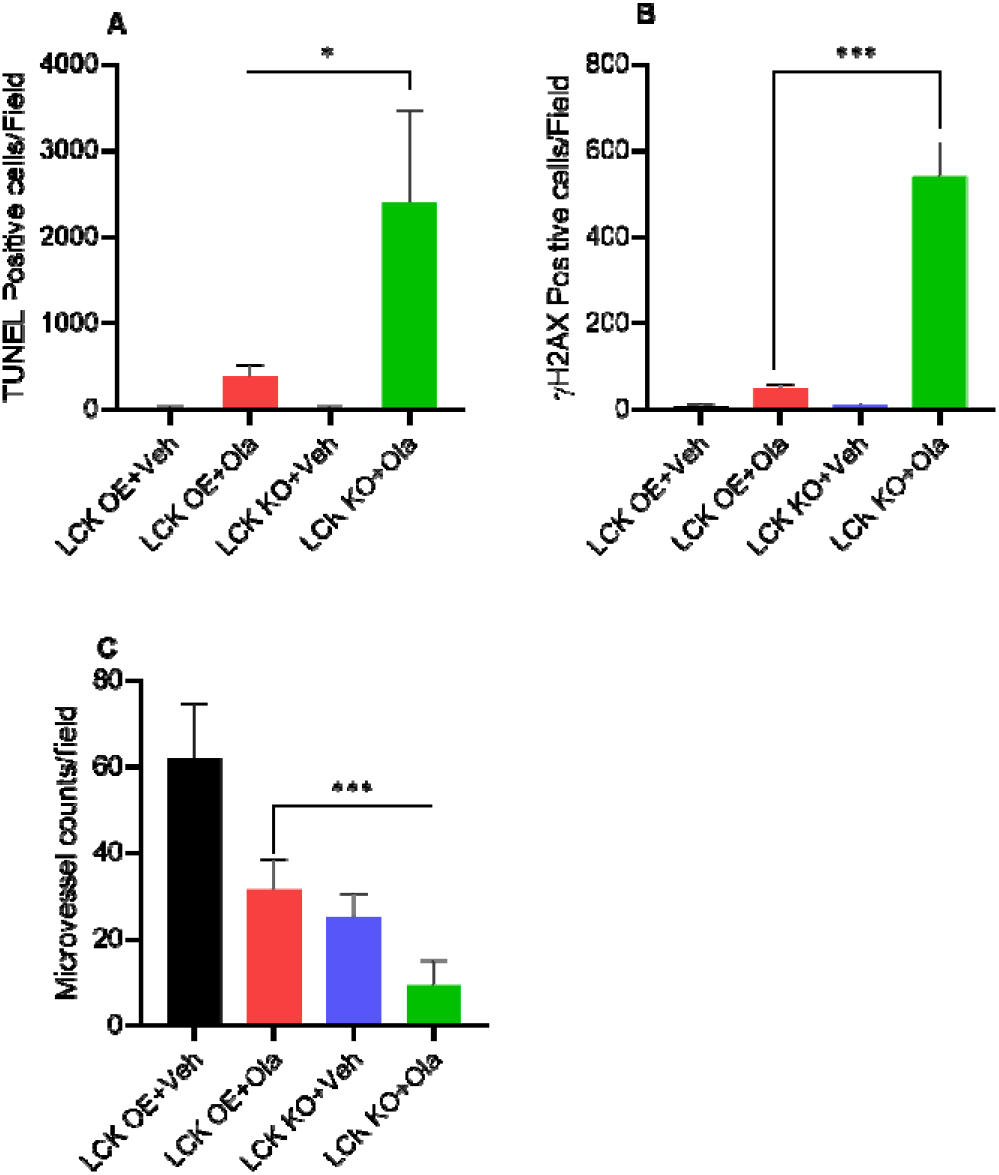
**(A)** TUNEL assay to detect DNA fragmentation in tumor tissue sections. TUNEL positive cells were counted from five images and plotted in graph (Main fig. 8D). **(B)** IHC staining of □H2AX of tumor sections from different groups. γH2AX positive cells were counted from five image and plotted in graph (Main fig. 8E).). **(C)** CD31 expression (Indicator of microvessel density and growth) of tumor sections from different group of mice. Microvessel density was counted from five images and plotted in graph (Main fig. 8F). Images are representative of two tumors from each cohort. We quantified the staining from 5 fields from each mouse. Images were captured at 20X magnification.

